# AntiSite: Unified paratope prediction for antibodies and nanobodies with or without structure

**DOI:** 10.64898/2026.08.24.745914

**Authors:** Angelos-Michail Papadopoulos, Federico Alvarez, Petros Daras

**Affiliations:** Information Technologies Institute, Centre for Research and Technology Hellas, Thessaloniki 57001, Greece; Universidad Politécnica de Madrid, Madrid 28040, Spain

**Keywords:** antibody, nanobody, paratope prediction, antibody structure prediction, deep learning, protein language models, molecular surface, antibody engineering

## Abstract

Accurate paratope prediction provides insights into antibody–antigen recognition while supporting antibody discovery and engineering. Existing predictors are fixed to a single input setting: sequence-based methods cannot use an antibody structure, structure-based methods require coordinates, and methods covering both combine separately trained predictors. We present AntiSite, an antigen-agnostic predictor whose sequence encoder, fusion block and prediction head are shared across both settings; structure-aware inference adds molecular-surface features from our previously developed ParaSurf model, used here as a frozen, separately trained structural encoder. Structural-modality dropout enables both inference settings without retraining. Across five antibody–antigen benchmarks, structure-aware AntiSite achieved the highest mean PR-AUC among all evaluated methods, while sequence-only PR-AUC was generally lower than that of the strongest dedicated antigen-agnostic sequence baseline. AntiSite also demonstrated zero-shot generalization to nanobodies without additional training. We further evaluated structures from recent antibody-specific and general protein folding methods, including Ibex, Protenix-v1, Boltz-2 and Chai-1, in antibody-only or antigen-aware co-folding settings. Their benefit was associated with structural accuracy, with the most accurate inputs yielding performance close to that obtained with experimental structures. AntiSite thus provides a shared predictor for antibody and nanobody binding sites, usable from sequence alone and refined when reliable structural information is available. Code and trained checkpoints are openly available.

## 1. Introduction

Therapeutic antibodies rely on highly specific interactions with their target antigens and have become a major class of modern biologic drugs[1]. These interactions are mediated by the paratope, the set of antibody residues that directly contact the antigen. The paratope is usually concentrated in the complementarity-determining regions (CDRs), although framework residues can also participate[2]. Predicted paratopes could inform residue selection for mutational analysis or serve as candidate constraints for antibody–antigen docking[3]. High-resolution antibody–antigen complex structures reveal these contacts directly, but are costly and time-consuming to determine experimentally and remain unavailable for many antibodies of interest. Computational prediction can therefore provide binding-site information from the antibody sequence or structure alone.

Sequence-based paratope predictors range from convolutional and recurrent networks, such as Parapred and ParaDeep, to protein language model approaches such as ParaAntiProt and Paraplume[2,4–6]. These methods operate directly on antibody sequences. ParaEM additionally incorporates the matching antigen sequence using an expectation–maximization framework[7]. Structure-based predictors, including Paragraph and our previously developed ParaSurf model, instead use antibody structures to capture local geometry and molecular-surface properties [8,9]. PECAN additionally uses the antigen structure to model interactions between the two binding partners[10]. Hybrid methods use both sequence and structural information. MIPE combines sequence and structural information from both the antibody and antigen to predict paratopes and epitopes[11]. Paraplume-G, for example, uses the structure-based model Paragraph for the CDRs and nearby residues, and the sequence-based model Para-plume for the rest of the antibody[2]. The two models are trained separately.

Existing predictors typically operate in a fixed input setting (Supplementary Table S4). Sequence-based methods do not accept antibody structural input, whereas structure-dependent methods require coordinates. Combinations such as Paraplume-G cover both settings using predictions from separately trained models. AntiSite instead trains a shared downstream predictor (its sequence encoder, fusion block and final prediction head) to operate in both settings (Figure 1). Structure-aware inference additionally uses a frozen ParaSurf encoder pretrained for supervised paratope prediction. Its internal features enter the shared AntiSite predictor, rather than combining the two models’ final scores. Neither inference mode requires antigen input.

**Figure 1.**
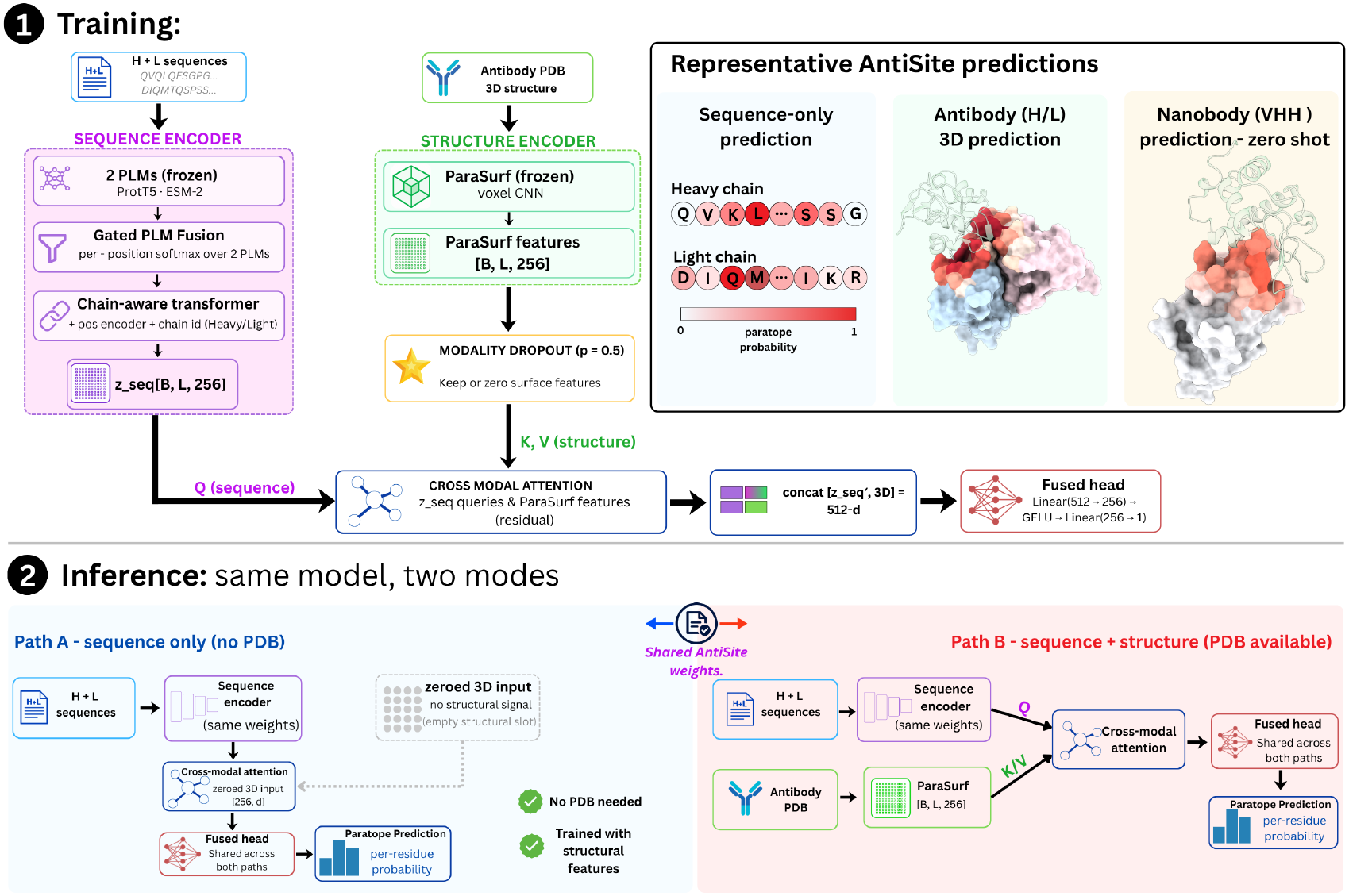
Overview of AntiSite. (1) **Training:** Frozen ProtT5 and ESM-2 embeddings pass through gated fusion and a chain-aware Transformer. A separately paratope-trained, frozen ParaSurf encoder provides optional molecular-surface features from experimental or predicted antibody structures. Cross-modal attention combines sequence and surface representations before a shared prediction head. Structural-modality dropout (*p* = 0.5) zeros surface features in half the training batches. (2) **Inference:** Sequence-only prediction (Path A) uses zero structural features; structure-aware prediction (Path B) uses ParaSurf features. Both share the AntiSite prediction checkpoint and output per-residue paratope probabilities. Examples show sequence-only prediction, a conventional antibody (1BVK) and zero-shot VHH prediction (1JTO). Pale-green antigens provide context only and are never model inputs.

Advances in structure prediction make this flexibility increasingly relevant. Models such as ABodyBuilder3 and Ibex predict antibody structures without antigen input[12,13]. Ibex can model both unbound (apo) and bound-like (holo) conformations. General models such as Protenix-v1 can also predict an antibody structure without its antigen[14]. However, variable-loop conformations, particularly CDR-H3, remain challenging to predict accurately[15].

At the same time, AlphaFold 3 and models such as Boltz-2, Chai-1, Protenix-v1 and ESMFold2 can jointly predict antibody–antigen complexes[14,16–19]. Contacts in these predicted complexes provide another way to estimate the paratope. These developments raise two practical questions: how much does a predicted antibody structure improve paratope prediction, and does including the antigen during structure prediction provide a better estimate of the binding site? Our comparisons focus on paratope prediction; complex-prediction methods also predict the antigen epitope and the relative binding pose.

Reliable evaluation of paratope predictors also requires testing on antibodies distinct from those used for training. Studies of antibody binding and affinity prediction have highlighted challenges in generalizing to unseen sequences and complexes [20–22]. Related work on nanobody non-specific binding has also shown how sequence clustering can influence reported predictive performance [23]. Together, these findings from related tasks motivate controlling sequence similarity between training and test antibodies, alongside evaluation on recently determined antibody–antigen complexes.

Transfer across antibody formats presents a further challenge. Nanobodies recognize antigens through a single heavy-chain variable domain (VHH), without a paired light chain. Recent work such as AbNatiV2 explicitly models sequence properties of conventional antibodies and nanobodies for computational design[24]. Whether a paratope predictor trained on paired antibodies can locate nanobody binding sites without additional training is a separate question. Evaluating the same trained model on genuine VHH–antigen complexes tests whether its learned representations transfer across these distinct antibody formats.

AntiSite combines ProtT5 and ESM-2 protein language model representations[25, 26] with molecular-surface features from frozen ParaSurf. Structural-modality dropout enables this flexibility, allowing molecular-surface information to complement sequence representations when an antibody structure is available. Even in sequence-only mode, AntiSite uses weights shaped by structural information during training, without requiring an antibody structure at inference. We evaluate both modes on antibody benchmarks and assess zero-shot transfer of the paired-antibody-trained model to nanobodies. A second contribution is a systematic evaluation of antibody-only folding and antigen-aware co-folding for paratope prediction. We test whether these predicted structures improve AntiSite, compare its predictions with direct co-folded interfaces, and examine how structural accuracy and predictor confidence relate to performance. Our results show that the shared AntiSite predictor supports sequence-only prediction and, with experimental antibody structures, achieves the highest mean PR-AUC among all evaluated methods across five antibody benchmarks. It also generalizes to nanobodies without additional training. We further show that the benefit of predicted structures depends on their accuracy. Together, AntiSite and these evaluations give immunologists a tool for locating candidate antibody and nanobody binding sites, along with evidence to guide the use of predicted structures.

## 2. Results

### 2.1. A shared predictor supports sequence-only and structure-aware prediction

AntiSite predicts the paratope from antibody sequence, with the option to add an experimental or predicted antibody structure (Figure 1). We refer to the latter mode as structure-aware AntiSite, labeled AntiSite (3D) in tables. Frozen ProtT5 and ESM-2 representations are combined by a learned gate and a chain-aware Transformer. When a structure is provided, the frozen ParaSurf encoder provides per-residue molecular-surface features[9]. Cross-modal attention combines the sequence and surface representations, and a shared head assigns paratope probabilities. Both modes use the same AntiSite sequence encoder, fusion block and prediction head, with no antigen input. ParaSurf is an additional task-trained network in structure-aware mode; its classification head is unused.

Modality dropout makes the structural input optional by replacing its features with zeros in half of the training batches. In a single-seed ablation on Paragraph-expanded, omitting this step reduced sequence-only PR-AUC from 0.736 to 0.496, while structure-aware PR-AUC changed by less than 0.001 (Supplementary Table S6B). Exposure to both input settings during training preserved sequence-only performance without reducing structure-aware accuracy.

A surface-only prediction head trained on frozen ParaSurf features reached mean PR-AUC 0.789 on Paragraph-expanded and 0.768 on PECAN across five seeds, below full AntiSite by 0.025 and 0.033, respectively. Sequence–surface concatenation recovered most of this improvement. Adding one cross-modal attention block provided smaller additional structure-aware gains of 0.002 on Paragraph-expanded and 0.004 on PECAN, with no consistent benefit in sequence-only mode (Supplementary Table S5). Full ablations are reported in Supplementary Section S3.

Figure 2 illustrates structure-aware predictions for three conventional antibodies and three VHHs using the same checkpoint trained on paired antibodies. Experimental antigens are shown only to locate the observed interfaces and were not used at all as input to AntiSite.

**Figure 2.**
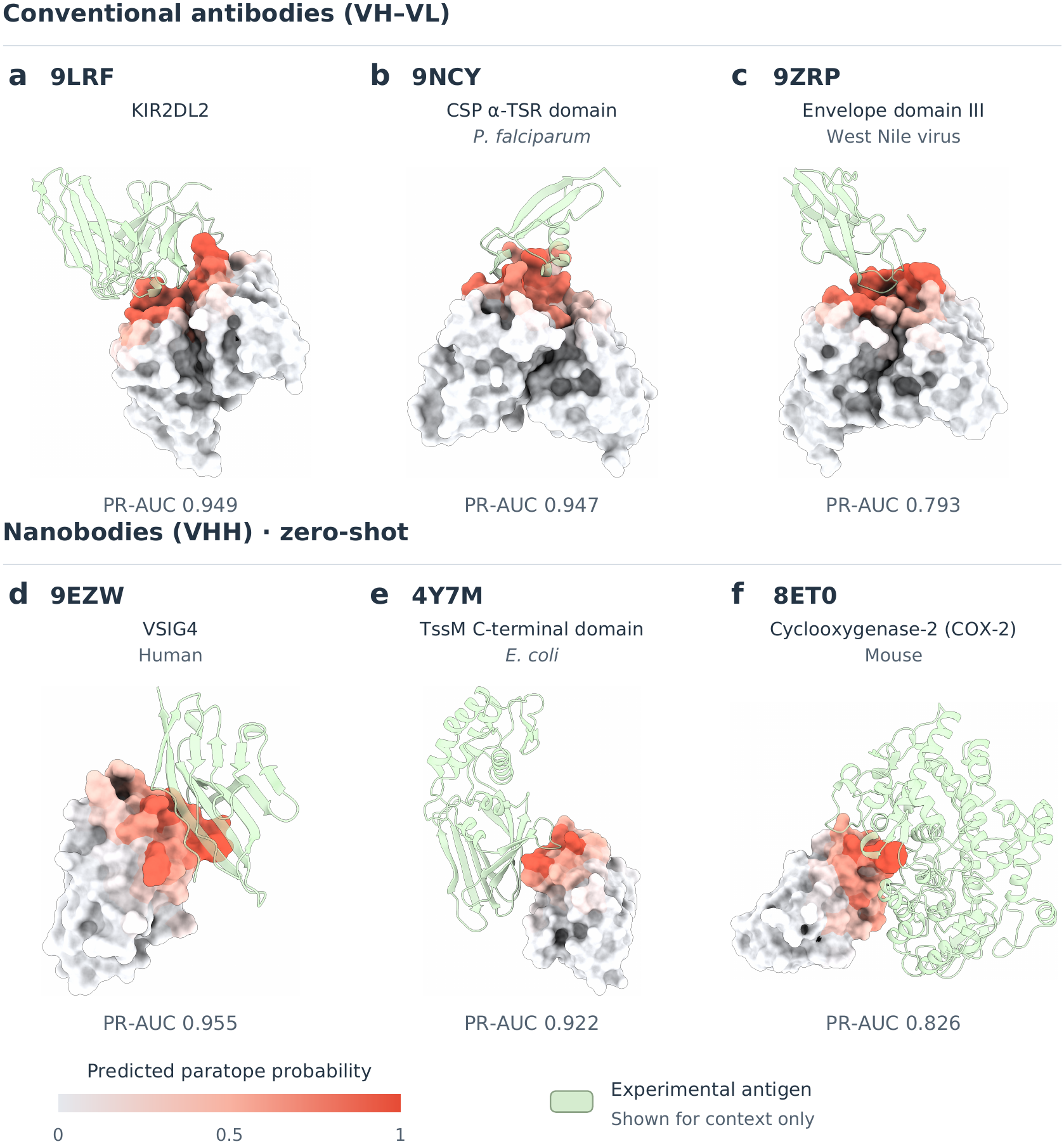
Structural examples of antibody and nanobody paratope prediction. Structure-aware predictions for conventional antibodies from SAbDab2 (a, 9LRF; b, 9NCY; c, 9ZRP) and zero-shot VHHs (d, 9EZW; e, 4Y7M; f, 8ET0), using experimental coordinates and the same paired-antibody-trained checkpoint. Surfaces range from pale gray to red with increasing paratope probability. Labels give PDB identifiers, antigens and PR-AUC over all scored residues. Each row contains two strong examples and one near its cohort median (c, f). Pale-green antigens show experimental binding context only and are not model inputs. Views are independently scaled; visualization details are in Supplementary Section S6.

### 2.2. Comparison with existing paratope predictors

We evaluated AntiSite on PECAN, Paragraph-expanded, MIPE and AACDB using their benchmark-specific training protocols, and tested the released Paragraphexpanded checkpoint on SAbDab2 without retraining[8,10,11,27,28]. Paragraph-expanded denotes the larger SAbDab benchmark of 1,086 antibody–antigen complexes introduced in the Paragraph study, before our preprocessing. Table 1 summarizes all five evaluations. The positional baseline assigns each residue the contact frequency observed at its sequence position in the training data. PR-AUC was the primary metric because contact residues are a minority of the antibody variable domain; ROC-AUC, F1 and MCC provide complementary measures.

**Table 1.**
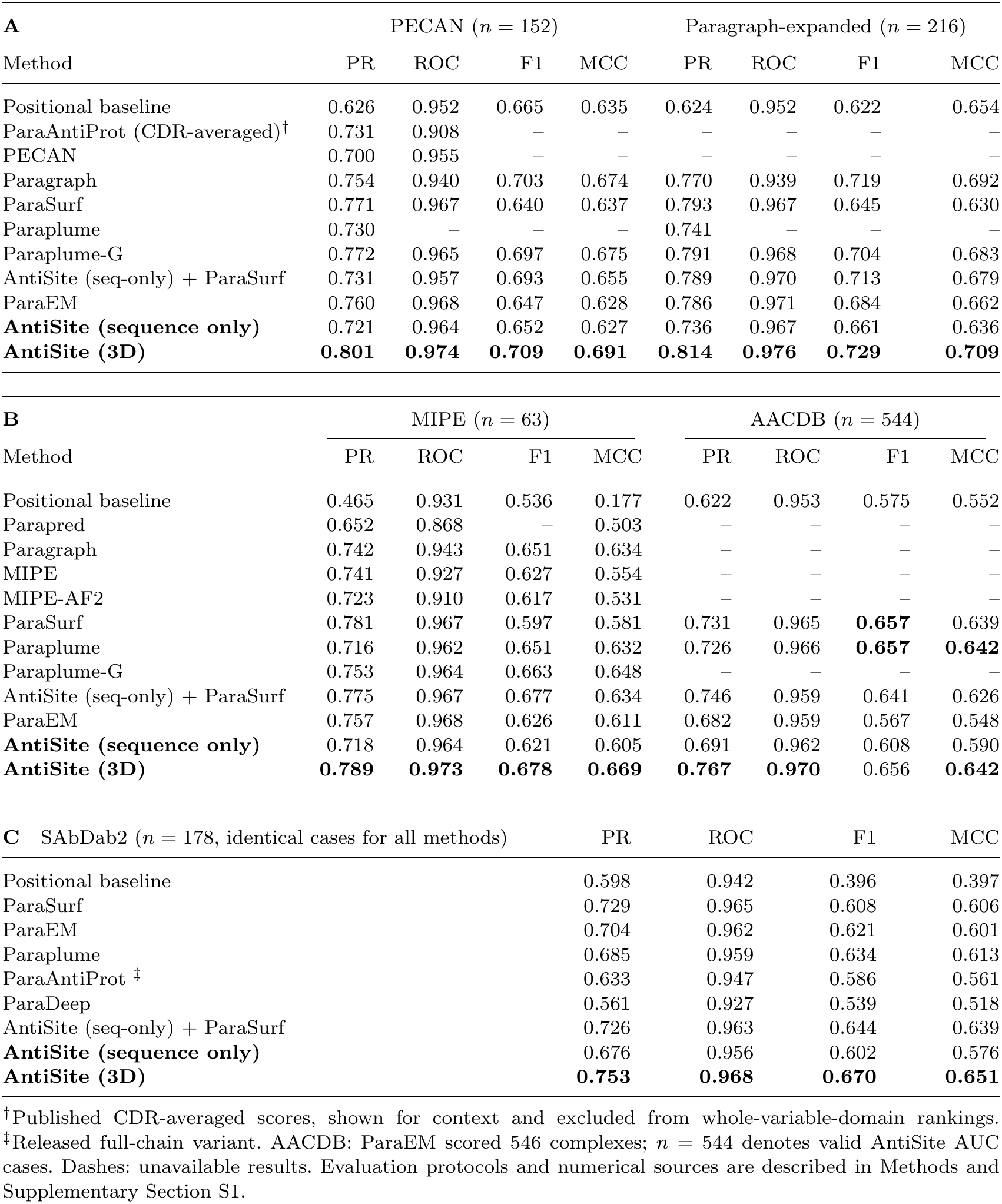
Comparison with existing paratope predictors across five antibody-antigen benchmarks. PR-AUC (PR) is the primary metric; ROC denotes ROC-AUC. Scores are per-complex means except the CDR-averaged entry. Bold marks the highest reported whole-variable-domain scores at the displayed precision. The two AntiSite modes use the same checkpoint within each evaluation.

With experimental antibody structures, structure-aware AntiSite achieved the highest PR-AUC and ROC-AUC among all evaluated methods across all five antibody benchmarks. PR-AUC was 0.801 on PECAN, 0.814 on Paragraph-expanded, 0.789 on MIPE, 0.767 on AACDB and 0.753 on SAbDab2. It exceeded Paraplume-G, which combines separately trained sequence and structure predictors, by 0.023–0.036 PR-AUC where comparisons were available, and also exceeded the published ParaEM values. Benchmark partitions, scoring conventions and numerical sources are described in Methods and Supplementary Section S1. Panel C reports the matched local SAbDab2 evaluation on identical antibody residues and contact labels.

On AACDB, AntiSite, ParaSurf, Paraplume and ParaEM were trained on the shared training partition and evaluated on its held-out test set. This followed the benchmark-specific training approach used for the other three datasets with published partitions. Structure-aware AntiSite achieved 0.767 PR-AUC, compared with 0.731 for ParaSurf and 0.682 for ParaEM. ParaEM additionally received antigen sequence; AntiSite used antibody inputs only. Baseline protocols and robustness analyses are provided in Supplementary Section S1.

We next evaluated the released Paragraph-expanded AntiSite checkpoint on a separate SAbDab2 test set, without retraining. Antibody-similarity filtering against the available training references and preprocessing checks yielded a final test set of 178 antibody–antigen complexes. The individual prediction methods were evaluated on this same filtered test set using identical antibody residues and contact labels. Structure-aware AntiSite again led PR-AUC (0.753), ahead of ParaSurf (0.729) and antigen-aware ParaEM (0.704), and achieved the highest means on the secondary metrics. The paired distributions show gains across individual complexes (Figure 3a): mean improvements over ParaSurf and ParaEM were 0.023 and 0.049, respectively, and remained significant after Holm adjustment[29] (*p*_adj_ = 5.8 *×* 10*^−^*^4^ and 2.8 *×* 10*^−^*^6^). Dataset construction, source evaluation conventions and full paired statistics are provided in Supplementary Section S1.

**Figure 3.**
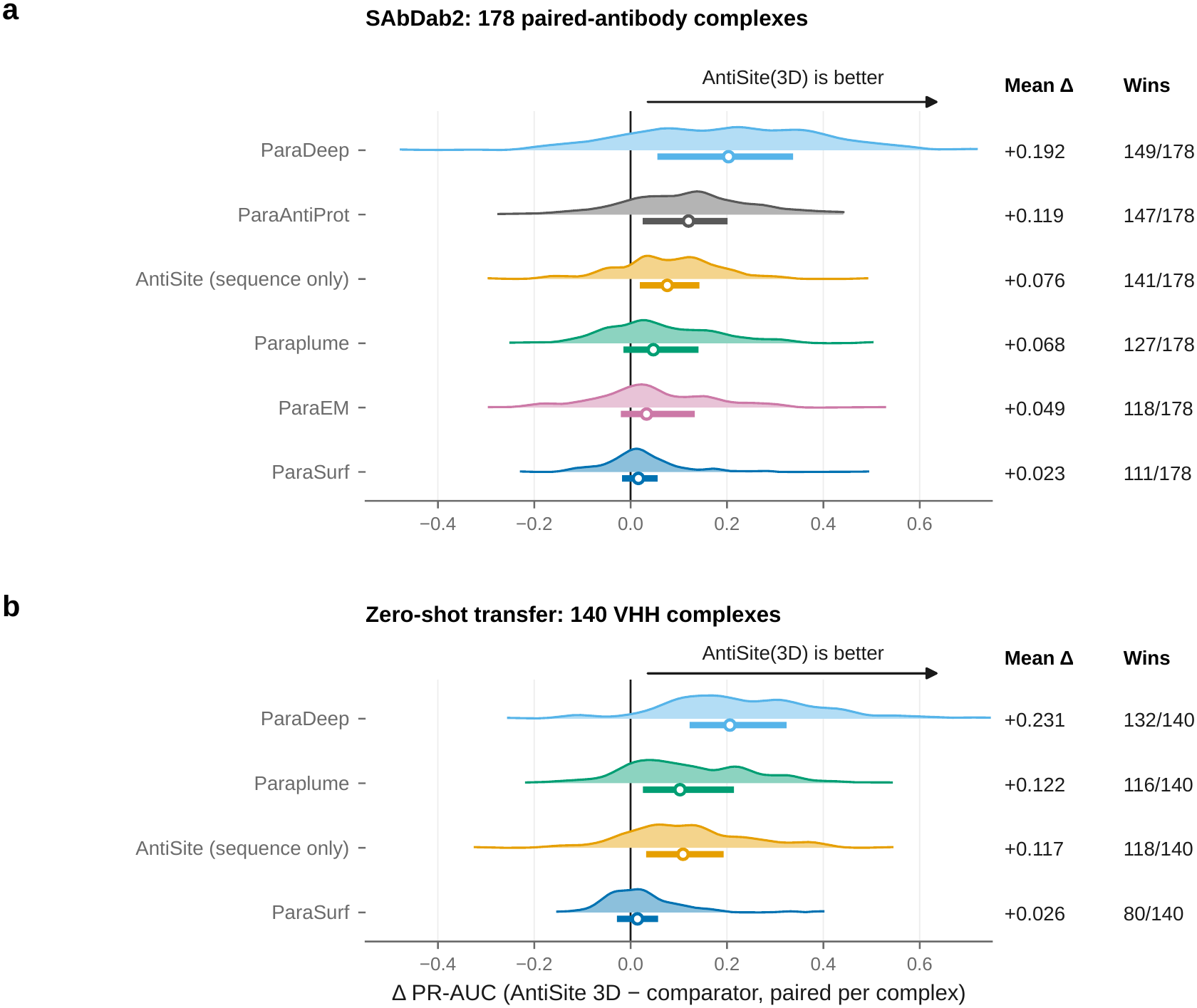
Paired improvements across antibody and nanobody complexes. Per-complex PR-AUC differences between structure-aware AntiSite and comparators on (a) 178 SAbDab2 and (b) 140 zero-shot VHH complexes. Right of zero favors AntiSite; left favors the comparator. Wider violin regions contain more complexes; white circles mark medians and horizontal bars the middle 50% of differences. Columns give mean differences and counts favoring AntiSite. Comparisons within each panel use identical complexes and the same paired-antibody-trained AntiSite checkpoint. Two-sided Wilcoxon tests with Holm adjustment within each panel are reported in Supplementary Table S3; baseline variants are described in Supplementary Section S1.

Sequence-only AntiSite had lower reported mean PR-AUC than Paraplume on four of the five evaluations: by 0.009 on PECAN, 0.005 on Paragraph-expanded, 0.035 on AACDB and 0.009 on SAbDab2. On MIPE, its PR-AUC was 0.002 higher. Both AntiSite modes achieved higher PR-AUC than the positional baseline on all five evaluations.

Combining the final residue scores from sequence-only AntiSite and standalone ParaSurf did not match AntiSite’s learned feature fusion (Table 1). The equal-weight ensemble also had lower reported PR-AUC than standalone ParaSurf on four of the five benchmarks; AACDB was the exception. Structure-aware AntiSite had higher mean scores than the ensemble on all four metrics across all five evaluations. On SAbDab2, the ensemble reached 0.726 PR-AUC, below structure-aware AntiSite (0.753) and ParaSurf alone (0.729).

### 2.3. Zero-shot generalization to nanobody paratopes

To test transfer across antibody formats, we applied the released paired-antibody-trained checkpoint to 140 genuine VHH–antigen complexes without additional training (Table 2). These are single-domain antibodies with complete paratopes, rather than conventional antibodies with their light chains removed. The cohort had no overlap with AntiSite or ParaSurf training data by PDB code or exact CDR-H3 sequence.

**Table 2.** Zero-shot paratope prediction on 140 nanobody–antigen complexes. Values are per-complex mean *±* SD. AntiSite uses the released paired-antibody-trained checkpoint without VHH training or fine-tuning. All methods are scored on the same VHH residues and contact labels; F1/MCC use 0.5. Bold values mark the highest means.

| Method | PR-AUC | ROC-AUC | F1 | MCC |
|---|---|---|---|---|
| ParaDeep | 0.538 ± 0.177 | 0.861 ± 0.097 | 0.521 ± 0.150 | 0.448 ± 0.168 |
| Paraplume | 0.647 ± 0.152 | 0.916 ± 0.074 | 0.578 ± 0.143 | 0.522 ± 0.160 |
| AntiSite (sequence only) | 0.651 ± 0.166 | 0.907 ± 0.084 | 0.534 ± 0.163 | 0.485 ± 0.182 |
| ParaSurf | 0.743 ± 0.187 | 0.932 ± 0.080 | 0.599 ± 0.181 | 0.557 ± 0.193 |
| AntiSite (3D) | **0.769** ± 0.169 | **0.941** ± 0.072 | **0.631** ± 0.196 | **0.591** ± 0.206 |

Structure-aware AntiSite achieved the highest means across all four metrics, including PR-AUC 0.769 compared with 0.743 for ParaSurf. Its paired gain was 0.026 over ParaSurf and 0.117 over sequence-only AntiSite; both remained significant after Holm adjustment (Figure 3b). The gain over ParaSurf also remained significant after averaging within the 104 antibody identities (*p*_adj_ = 0.012; Supplementary Table S3B). Sequence-only PR-AUC was 0.651, compared with 0.647 for Paraplume, although its ROC-AUC, F1 and MCC were lower. The same model therefore transferred to VHHs in both input settings, with structural information providing the larger gain.

Examples of these VHH predictions are shown in Figure 2d–f.

### 2.4. Structural input preferentially raises paratope probabilities

For the following structural analyses, we selected 87 antibody–antigen complexes released in the PDB on or after 1 February 2024. The final set comprised 30 complexes released in 2024, 43 in 2025 and 14 in 2026. This selection aimed to reduce known training-data overlap and provide a common basis for comparing antibody-only structure predictors, such as Ibex, with antigen-aware co-folding models, such as Boltz-2. The date cutoff followed ABodyBuilder3’s documented January 2024 training boundary[12]; additional filtering criteria, predictor-specific cutoffs and processing details are given in Supplementary Section S4.

Using this cohort, we first compared sequence-only and structure-aware predictions with experimental antibody structures. Adding experimental antibody structure increased probability by 0.110 on average at true paratope residues, while the mean change at other residues was *−*0.004 (Figure 4b,c). The orange curve is generally positive, while the gray curve stays closer to zero: structure mainly raises scores at actual contacts. Figure 4c shows the same pattern after grouping residues by their sequence-only probabilities: true contact residues generally lie above the diagonal, while other residues lie close to or below it. Antibody 9JBQ illustrates this effect: its PR-AUC increased from 0.626 to 0.898 (Figure 4a).

**Figure 4.**
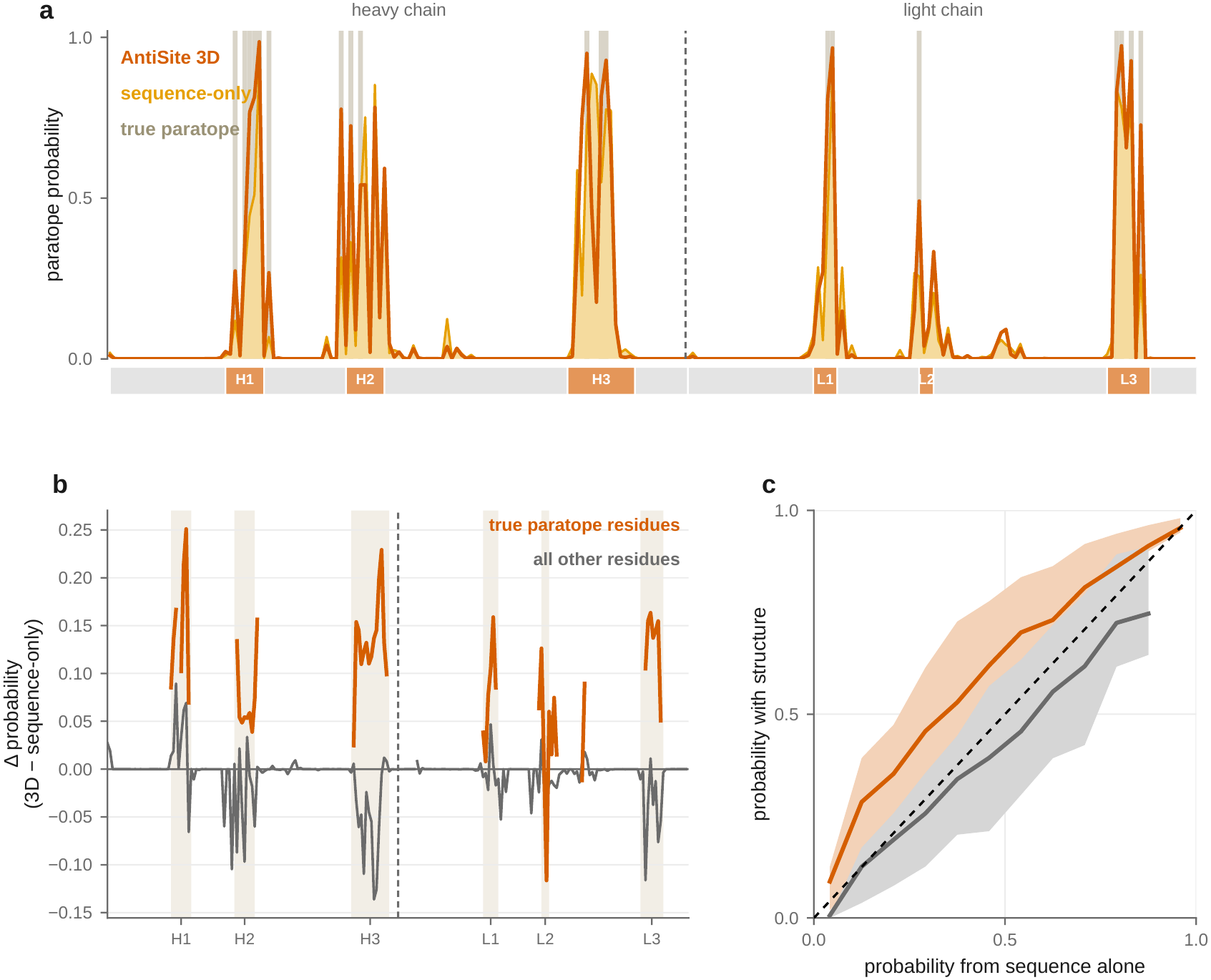
Residue-level changes when experimental antibody structure is provided. Both modes use the same checkpoint. (a) Predictions for 9JBQ: lighter, sequence-only; darker, structure-aware. Gray bands mark true binding residues, the dashed line separates chains and the strip marks CDRs. (b) Mean probability changes across 87 antibodies: structure-aware minus sequence-only. Above zero means an increase; below zero a decrease. Orange marks true contacts and gray other residues. (c) Structure-aware probabilities plotted against sequence-only probabilities for the same two groups. Above the black diagonal means increased probability; below means decreased probability. Curves show means and bands the middle 50% of scores. Contact labels are used only for analysis. Alignment and resampling details are in Supplementary Section S6.

The regional analysis on Paragraph-expanded also showed improvement within the CDRs: CDR-H3 PR-AUC increased from 0.824 in sequence-only mode to 0.902 with structure. In the matched framework comparison, structure-aware AntiSite also achieved higher PR-AUC than ParaSurf (0.616 versus 0.546), together with higher ROC-AUC. At threshold 0.5, AntiSite made fewer framework predictions with higher precision, whereas ParaSurf recovered more contacts (Supplementary Section S6).

### 2.5. Paratope prediction with antibody-only and co-folded structures

We first asked whether an antibody structure predicted from its heavy- and light-chain sequences alone could improve AntiSite predictions when no experimental antibody structure is available (Table 3A). We used the frozen AntiSite checkpoint trained on Paragraph-expanded throughout these analyses.

**Table 3.**
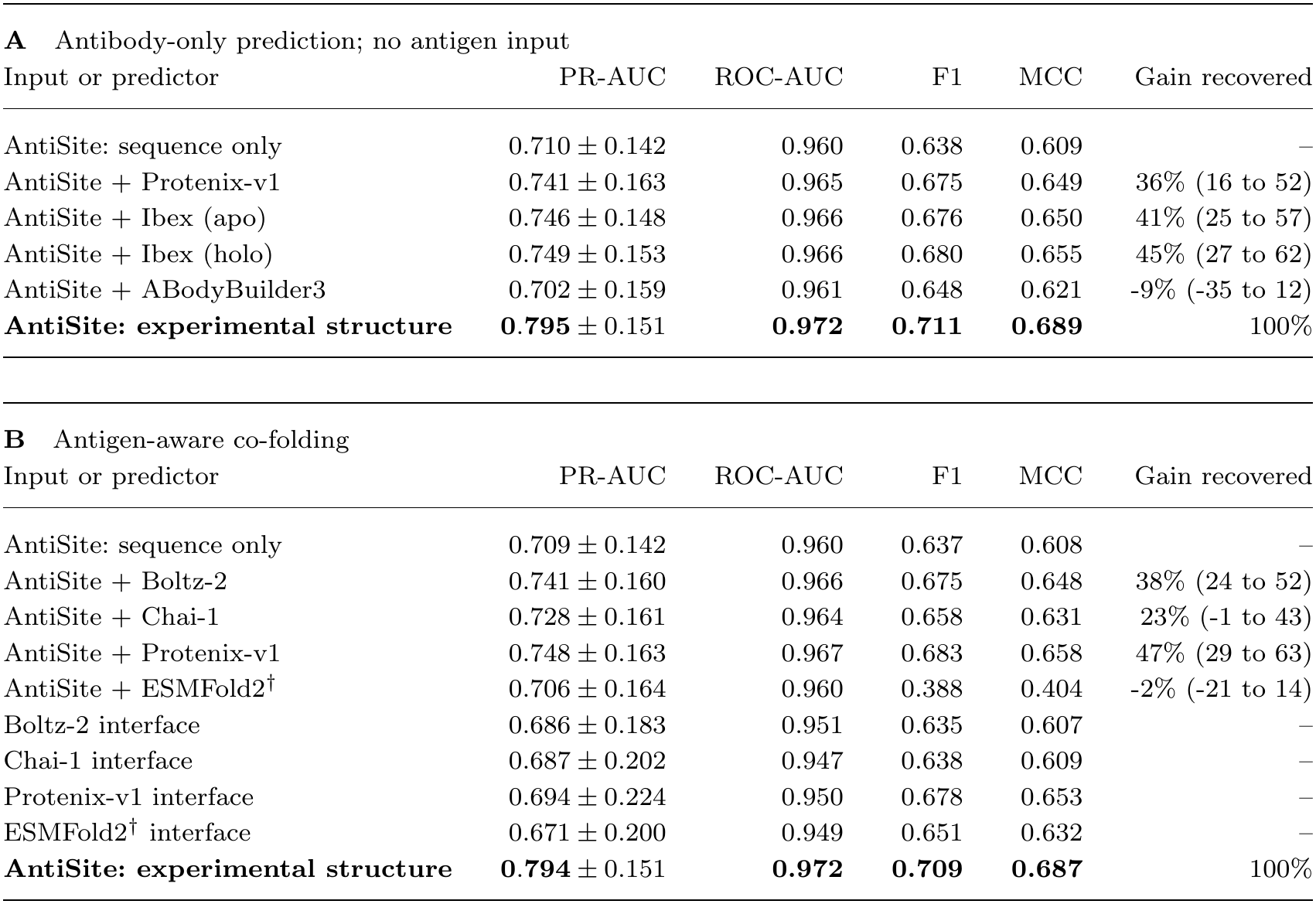
Paratope prediction using predicted antibody structures and direct co-folded interfaces. A: antibody-only prediction, 87 complexes. B: antigen-aware co-folding, 86 shared complexes, except ESMFold2 (*†*, 80). PR-AUC is mean *±* SD across complexes; other metrics are means. All AntiSite entries use the frozen checkpoint trained on Paragraph-expanded, without retraining.

| **A** Antibody-only prediction; no antigen input |  |  |  |  |  |
| Input or predictor | PR-AUC | ROC-AUC | F1 | MCC | Gain recovered |
|---|---|---|---|---|---|
| AntiSite: sequence only | 0.710 ± 0.142 | 0.960 | 0.638 | 0.609 | – |
| AntiSite + Protenix-v1 | 0.741 ± 0.163 | 0.965 | 0.675 | 0.649 | 36% (16 to 52) |
| AntiSite + Ibex (apo) | 0.746 ± 0.148 | 0.966 | 0.676 | 0.650 | 41% (25 to 57) |
| AntiSite + Ibex (holo) | 0.749 ± 0.153 | 0.966 | 0.680 | 0.655 | 45% (27 to 62) |
| AntiSite + ABodyBuilder3 | 0.702 ± 0.159 | 0.961 | 0.648 | 0.621 | -9% (-35 to 12) |
| **AntiSite: experimental structure** | **0.795** ± 0.151 | **0.972** | **0.711** | **0.689** | 100% |
| **B** Antigen-aware co-folding |  |  |  |  |  |
| Input or predictor | PR-AUC | ROC-AUC | F1 | MCC | Gain recovered |
| AntiSite: sequence only | 0.709 ± 0.142 | 0.960 | 0.637 | 0.608 | – |
| AntiSite + Boltz-2 | 0.741 ± 0.160 | 0.966 | 0.675 | 0.648 | 38% (24 to 52) |
| AntiSite + Chai-1 | 0.728 ± 0.161 | 0.964 | 0.658 | 0.631 | 23% (-1 to 43) |
| AntiSite + Protenix-v1 | 0.748 ± 0.163 | 0.967 | 0.683 | 0.658 | 47% (29 to 63) |
| AntiSite + ESMFold2<sup>†</sup> | 0.706 ± 0.164 | 0.960 | 0.388 | 0.404 | -2% (-21 to 14) |
| Boltz-2 interface | 0.686 ± 0.183 | 0.951 | 0.635 | 0.607 | – |
| Chai-1 interface | 0.687 ± 0.202 | 0.947 | 0.638 | 0.609 | – |
| Protenix-v1 interface | 0.694 ± 0.224 | 0.950 | 0.678 | 0.653 | – |
| ESMFold2<sup>†</sup> interface | 0.671 ± 0.200 | 0.949 | 0.651 | 0.632 | – |
| **AntiSite: experimental structure** | **0.794** ± 0.151 | **0.972** | **0.709** | **0.687** | 100% |

Antibody-only structures from Protenix-v1 and Ibex improved PR-AUC over sequence-only inference, recovering 36–45% of the gain provided by experimental structures. Ibex holo, which predicts a bound-like conformation without antigen input, gave the highest mean performance. Paired two-sided Wilcoxon tests against sequence-only inference gave unadjusted *p ≤* 0.003 and are reported descriptively; per-input results are given in Supplementary Table S10B. Predictions using experimental structures remained more accurate on average. ABodyBuilder3 gave no average improvement. Gains were uncertain in the smaller subset released after Ibex’s weights became available (Supplementary Section S4).

We then asked whether knowing the antigen during structure prediction gives AntiSite a better antibody structure (Table 3B). Boltz-2, Chai-1, Protenix-v1 and ESM-Fold2 predicted complexes from antibody and antigen sequences; only the extracted antibody coordinates were provided as AntiSite’s structural input. The common comparison uses 86 complexes because one Chai-1 prediction was unavailable. ESMFold2 is reported separately on 80 complexes because predictions were unavailable for seven of the 87 cases.

Antigen-aware co-folding provided no consistent additional benefit over antibody-only prediction. The best co-folded input, Protenix-v1, gave a similar mean PR-AUC to antibody-only Ibex holo on the common cohort (both 0.748; unadjusted paired *p* = 0.89). For Boltz-2, Chai-1 and Protenix-v1, applying AntiSite to their antibody structures gave higher mean PR-AUC than reading contacts directly from their predicted complexes. Protenix-v1’s direct-interface F1 and MCC were nevertheless close to AntiSite with predicted antibody inputs, so the advantage was clearest in residue ranking.

ESMFold2’s saved side-chain atom overlaps distorted its surface features and compressed AntiSite probabilities, particularly affecting F1 and MCC at threshold 0.5. Processing details are given in Supplementary Section S4. The confidence analysis below examines cases where direct co-folded interfaces performed better than their overall averages suggest.

### 2.6. Benefits from predicted structures track their accuracy

Structural accuracy helped explain why predicted inputs differed in usefulness. Across 608 predicted structures for the same 87 antibodies, CDR-H3 backbone RMSD below 1.5 Å was associated with a mean PR-AUC gain of 0.066 over sequence-only inference, and 86% of predictions improved performance (Figure 5a). Mean gain fell to 0.023 at 1.5–3 Å and to *−*0.004 at 3 Å or above. The lowest-error eighth of predictions recovered 94% of the experimental-structure gain (95% bootstrap interval 82–106%), showing that accurate predicted inputs can approach the benefit of experimental structures.

**Figure 5.**
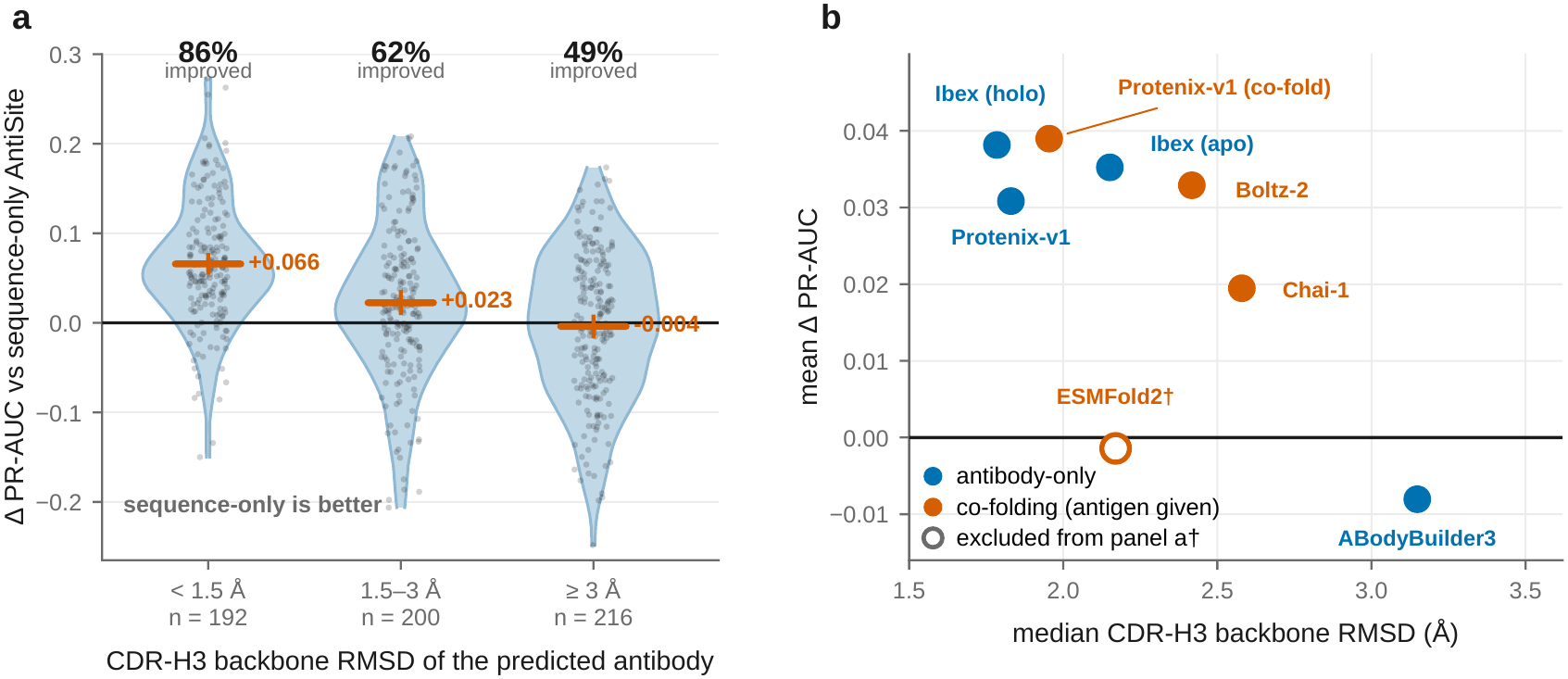
Structural accuracy is associated with the benefit of predicted antibody inputs. (a) PR-AUC gain over sequence-only prediction by CDR-H3 backbone RMSD. Points represent predicted structures; violins show their distributions. Orange markers show means and 95% bootstrap intervals; percentages give the fraction improved. The 608 predictions cover 87 antibodies; intervals resample predictions and are descriptive. (b) Mean gain against median RMSD for each condition: blue, antibody-only; orange, antigen-aware co-folding. ESMFold2 (open marker) appears only in b because of side-chain defects. RMSD requires an experimental reference and does not provide a prospective selection rule.

The association also held when comparing alternative structures of the same antibody: the mean within-complex Spearman correlation between CDR-H3 RMSD and PR-AUC gain was *−*0.38 (*p* = 1.9 *×* 10*^−^*^9^). Both antibody-only and co-folding predictions followed this trend (Figure 5b). ABodyBuilder3 had the largest median CDR-H3 error (3.15 Å), consistent with its lack of average benefit. ESMFold2 was excluded from this backbone-based analysis because its documented side-chain defects are not captured adequately by these RMSDs.

We compared the surface features from experimental structures with those from six predictors across eight prediction settings: ABodyBuilder3, Ibex (apo and holo), Protenix-v1 (antibody-only and co-folding), Boltz-2, Chai-1 and ESMFold2 (Figure 6). In panel a, each dot represents one residue’s ParaSurf features, summarized in a two-dimensional map. Most sources overlap, whereas ESMFold2 forms a partly distinct region.

**Figure 6.**
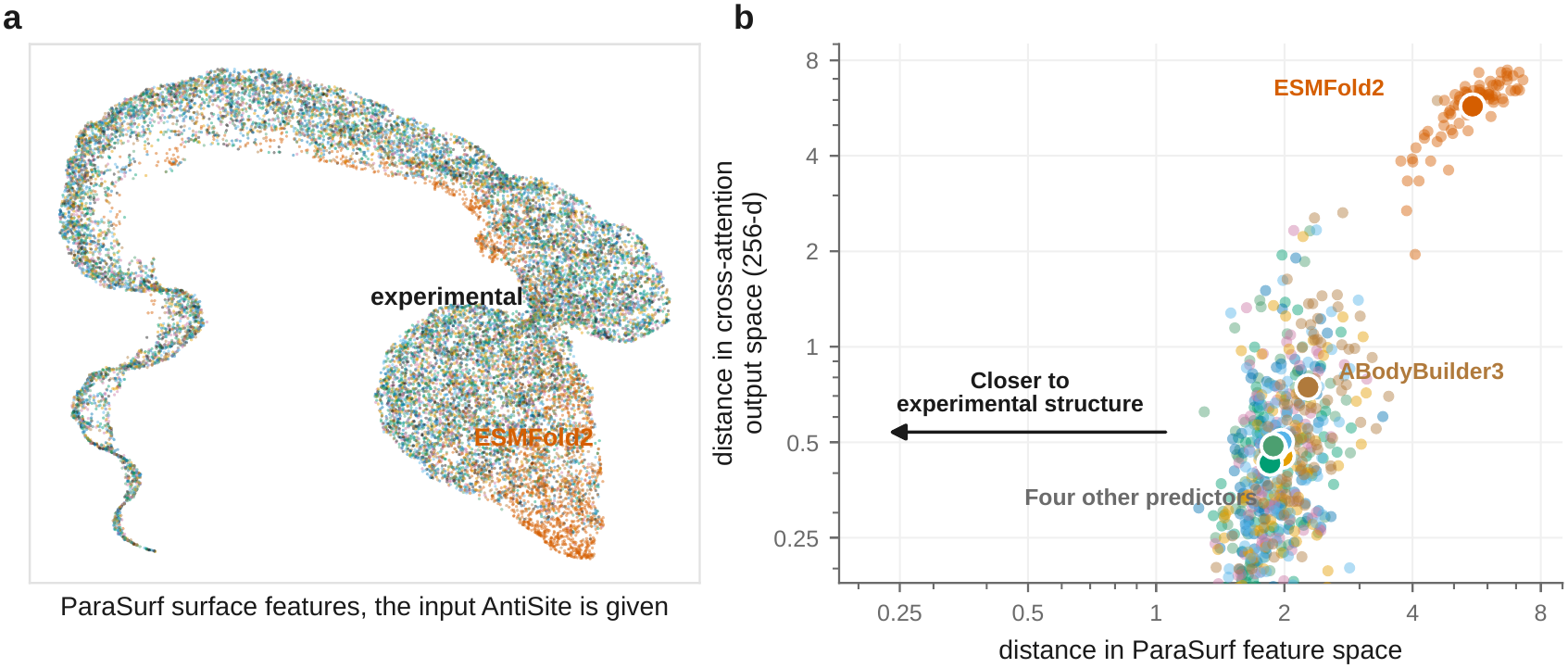
How predicted structures change the structural features used by AntiSite. Six predictors cover eight settings on the 87-antibody cohort; Ibex and Protenix-v1 each contribute two settings. (a) Each dot represents a residue’s ParaSurf features in a UMAP map, not its physical coordinates. Colors identify experimental and predicted inputs; most overlap, while ESMFold2 partly separates. (b) Each dot compares a predicted structure with its experimental reference: horizontal, ParaSurf-feature distance; vertical, cross-attention-output distance. Bottom-left points are closer to the reference in both spaces; large circles mark condition means. Both axes are logarithmic, with different, non-comparable scales. The four other predictors are Ibex, Protenix-v1, Boltz-2 and Chai-1. ESMFold2 is most displaced. The classifier also receives original surface features. Sampling and distance calculations are in Supplementary Section S6.

Panel b compares each predicted structure with its experimental reference. The horizontal axis shows the difference in ParaSurf features; the vertical axis shows the difference after cross-modal attention combines surface and sequence information. Points closer to the bottom left are closer to the experimental reference in both representations; large circles mark the average for each setting. ESMFold2 lies far toward the upper right, consistent with its documented side-chain problems. The two axes measure different learned representations and have different scales.

### 2.7. Predictor confidence indicates the expected benefit of structural information

We next examined whether the structure predictors’ own confidence scores could help identify reliable paratope predictions when no experimental antibody structure is available. Figure 7a compares predictions read directly from co-folded interfaces with Anti-Site predictions using antibody-only Ibex holo structures. Each point is one predicted complex; moving right means higher interface confidence (ipTM). The vertical axis shows the PR-AUC difference: above the horizontal zero line, the co-folded interface performs better; below it, AntiSite performs better. The thick black curve shows the average difference among predictions with similar confidence, and the gray band shows uncertainty around that average. Its upward trend indicates that co-folded interfaces become more competitive as confidence increases. Among high-confidence predictions from Boltz-2, Chai-1 and Protenix-v1 (ipTM *≥* 0.8; 50 of 260 predictions), mean PR-AUC was 0.818 for direct interfaces and 0.826 for AntiSite (*p* = 0.87). At lower confidence, direct interfaces remained less accurate on average. In model-specific high-confidence subsets, Chai-1, Protenix-v1 and ESMFold2 interfaces had higher mean PR-AUC than AntiSite, whereas Boltz-2 remained lower.

**Figure 7.**
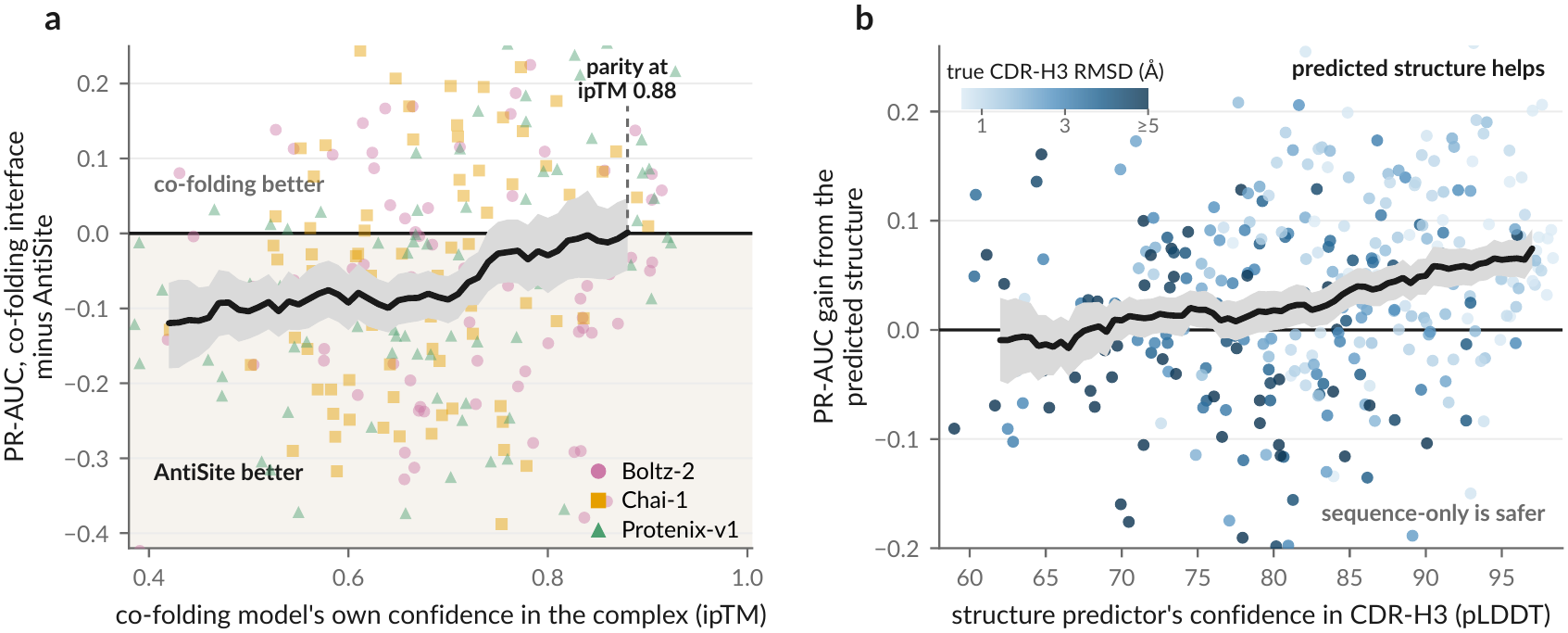
How structure-prediction confidence relates to paratope performance. (a) Direct-interface PR-AUC minus AntiSite PR-AUC with Ibex holo inputs, against ipTM, for 260 Boltz-2, Chai-1 and Protenix-v1 complexes. Above zero favors direct interfaces; below favors AntiSite. (b) AntiSite’s gain from structural input against CDR-H3 pLDDT for 347 predictions from antibody-only Protenix-v1 and the three co-folding models; color indicates CDR-H3 RMSD to the experimental reference. Above zero favors structural input; below favors sequence-only inference. Black curves show sliding-window means; gray bands are pointwise 95% bootstrap intervals. Repeated antibodies make these pooled summaries descriptive. The ipTM 0.88 crossing is a fitted trend, not a validated cutoff. ESMFold2 is excluded; model-specific results are in Supplementary Section S5.

Figure 7b examines whether providing a predicted antibody structure improves AntiSite over sequence alone. Each point is one predicted structure, with confidence in its CDR-H3 loop (pLDDT) on the horizontal axis and the PR-AUC gain from adding structure on the vertical axis. Above zero, structural input helps; below zero, sequence-only inference performs better. The thick black curve again shows the average gain among predictions with similar confidence, with uncertainty shown by the gray band. Its upward trend indicates that higher-confidence structures provide larger gains on average. Across 347 structures, higher CDR-H3 pLDDT was associated with lower structural error (*ρ* = *−*0.61). Scores of at least 90 were associated with a mean PR-AUC gain of 0.062, whereas scores below 70 gave no average benefit. However, using 70 as a cutoff for switching to sequence-only inference changed each predictor’s mean PR-AUC by only *−*0.004 to +0.009. We therefore recommend considering structures from antibody-only predictors that improved paratope prediction in this benchmark, using confidence as context rather than an automatic pLDDT-70 switch between modes. Detailed confidence bins and model-specific comparisons are provided in Supplementary Section S5, Table S15.

### 2.8. PLM comparisons support using ProtT5 and ESM-2 in AntiSite

Under AntiSite’s shared architecture and training settings, the general protein language model pair ProtT5 and ESM-2 gave higher mean PR-AUC than the tested antibody-specific AbLang2/IgT5 pair[30,31]. This held across five seeds on PECAN and Paragraph-expanded in both sequence-only and structure-aware modes; the general pair also outperformed the four-encoder stack containing all four models. On MIPE, the general pair and four-encoder stack performed similarly. These comparisons support the ProtT5/ESM-2 design choice within the evaluated training configuration (Supplementary Section S3, Table S7).

AntiSite also learned a pronounced regional preference without receiving CDR or framework annotations: ProtT5 received higher weights in CDRs, whereas ESM-2 received higher weights in frameworks (Figure 8). The gate mixes the two PLMs’ features at each antibody residue. In panels a and b, orange indicates the ProtT5 weight and blue the complementary ESM-2 weight. These are feature-mixing weights, not paratope probabilities. Panel a shows the typical pattern across 87 antibodies, with heavy chains on the left and light chains on the right. The black curve follows the median ProtT5 weight: values near 1 favor ProtT5, while values near 0 favor ESM-2. The dashed line at 0.5 marks equal weighting, the white band covers the middle 50% of weights across antibodies, and the strips identify CDRs and framework regions.

**Figure 8.**
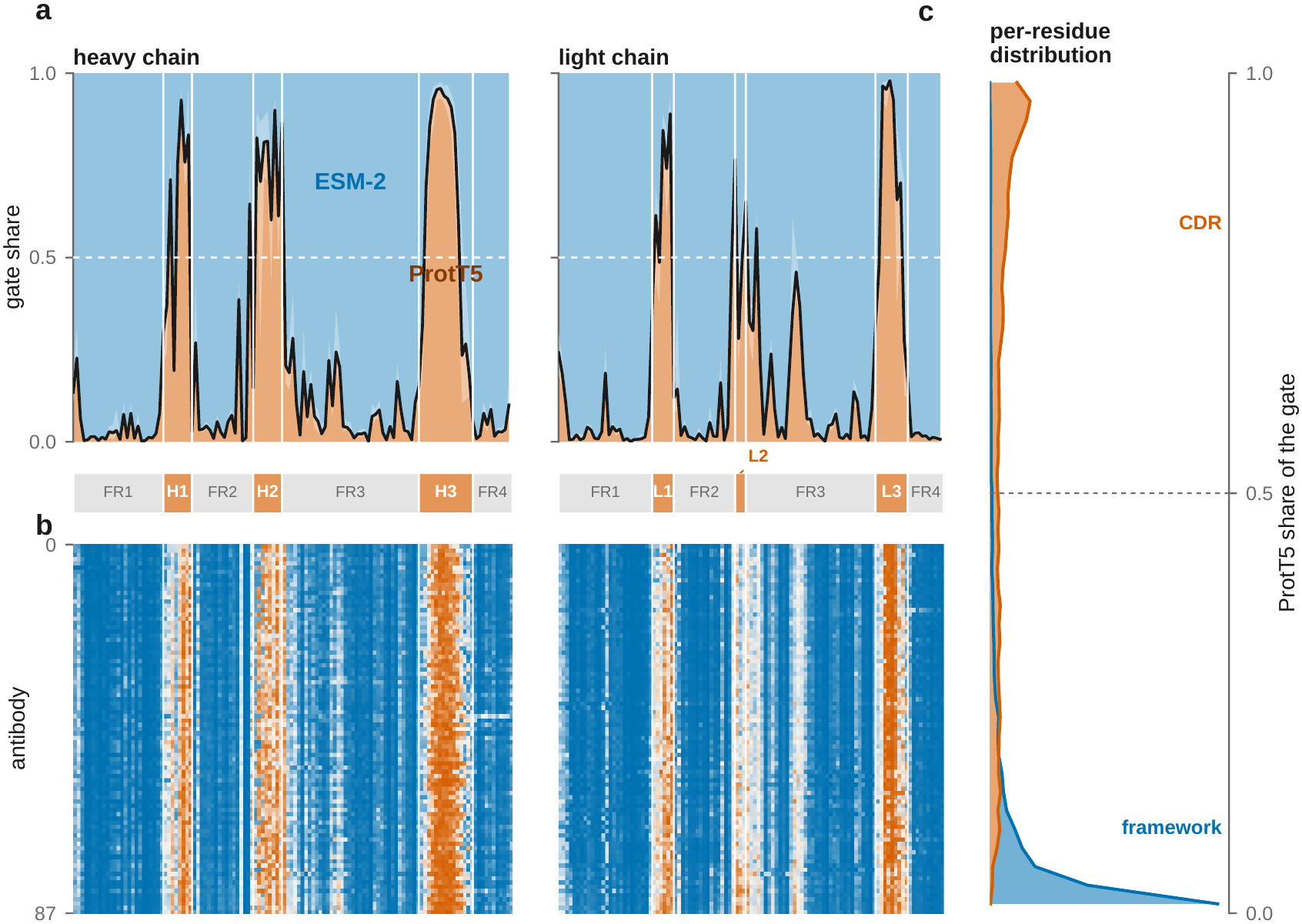
Learned regional allocation of protein language model representations. (a) Median heavy- and light-chain gate profiles across 87 antibodies: orange, ProtT5; blue, ESM-2. The black curve shows median ProtT5 share and the white band its interquartile range; the dashed line marks equal weighting. (b) Individual antibody profiles, one per row. (c) ProtT5-share distributions for CDRs (orange) and frameworks (blue), with medians 0.72 and 0.03. CDR annotations are used only for analysis. Gate weights describe feature mixing, not feature importance. Profile construction and sorting are described in Supplementary Section S6.

Panel b shows individual antibodies, one per row within each chain block; heavy- and light-chain rows are sorted separately. Orange vertical bands indicate that many antibodies give ProtT5 higher weights in similar CDR regions. Panel c summarizes the weight distributions for CDR residues (orange) and framework residues (blue). Its vertical axis again shows the ProtT5 weight; wider parts indicate more common weights within each group. The median ProtT5 weight was 0.72 in CDRs and 0.03 in frameworks, showing generally higher ProtT5 weighting in CDRs and higher ESM-2 weighting in frameworks. The gate learned this allocation through the paratope-prediction objective, without a predefined regional weighting rule. Gate weights describe the learned mixing rule; the predictive benefit of the encoders is assessed by the separate full-model ablations.

Figure 9 shows how the same antibody residues are represented at different stages of AntiSite. Each dot is one of 20,037 residues from 87 antibodies, shown in a two-dimensional UMAP map of its numerical features. Nearby points generally have similar features. The columns show frozen ProtT5, ESM-2 and ParaSurf inputs, followed by AntiSite’s sequence-encoder output and its output after surface attention. Within each column, both rows show the same points: the top marks experimentally observed contacts in orange, while the bottom distinguishes CDRs and frameworks.

**Figure 9.**
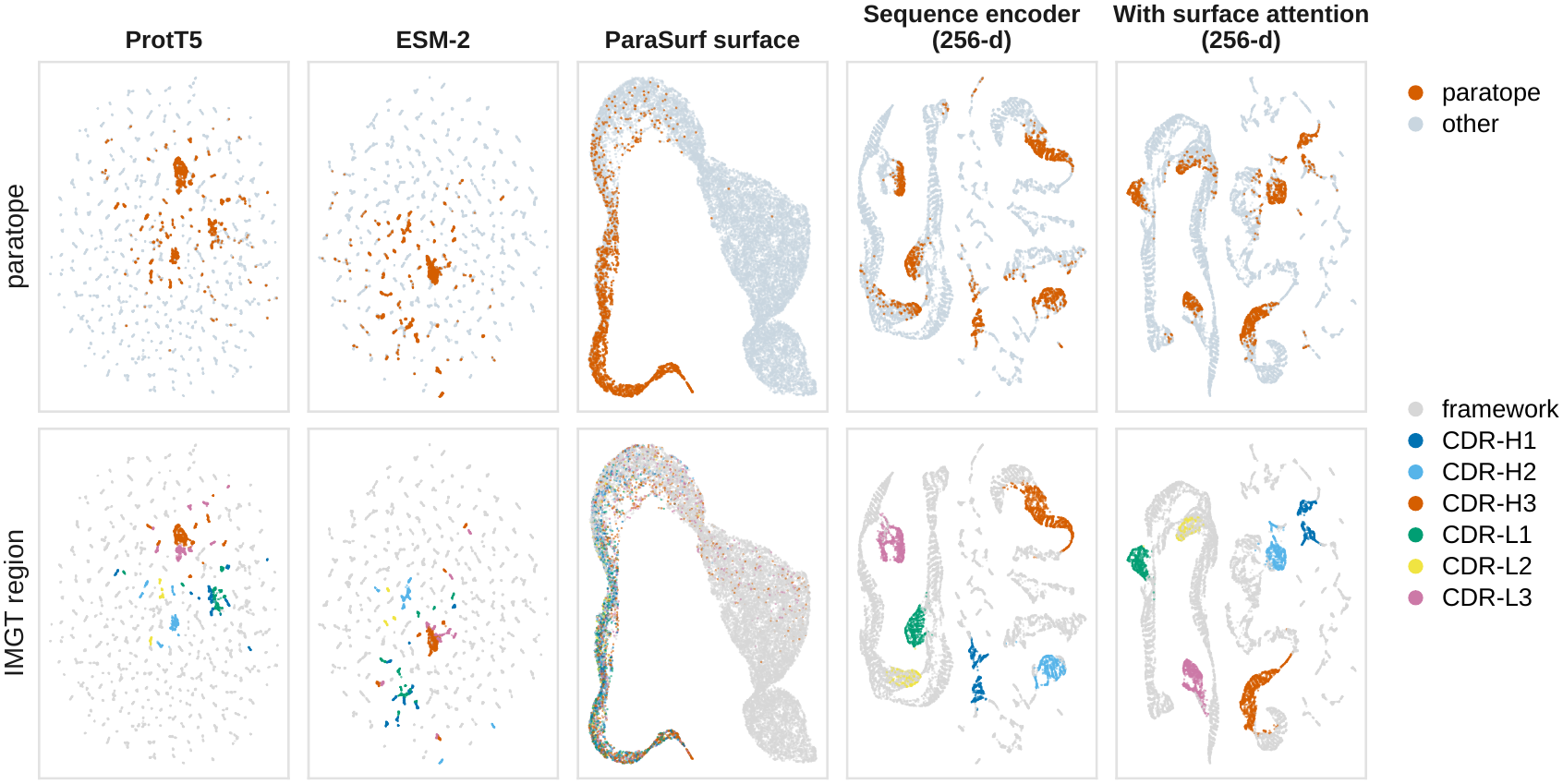
Antibody residue features at different stages of AntiSite. Each map shows the same 20,037 residues from 87 antibodies. Columns show ProtT5, ESM-2 and ParaSurf inputs, then sequence-encoder and surface-attention outputs. The sequence-encoder column precedes cross-modal attention. Top: contacts orange, other residues pale blue. Bottom: six IMGT CDRs colored separately, frameworks gray. Nearby points generally represent similar numerical features. Both rows share coordinates within each column; maps are fitted independently, so positions and distances cannot be compared across columns. CDR labels are applied only for analysis. These intermediate representations do not measure predictive accuracy; full-model results appear in Table 1. Projection settings and the relationship to inference outputs are described in Supplementary Section S6.

First, in the ParaSurf column, many contact residues occupy a common part of the map, where residues from different CDRs mix. This is consistent with shared surface features associated with antigen contact across different CDR loops.

Second, the last two columns show distinct CDR-associated groups, with actual contacts concentrated within parts of those groups. No CDR-category labels were provided to AntiSite during training; chain identity and residue position were explicit model inputs. The maps were fitted from features alone and colored afterward. Each projection was fitted independently, so these views cannot rank predictive accuracy or quantify the benefit of attention; the full-model comparisons in Table 1 establish performance.

### 2.9. Computational cost of the two inference modes

With all models resident in memory on an NVIDIA RTX 3090 workstation, sequence-only prediction took approximately 0.1 s per input. Structure-aware prediction averaged 13.2 s for Fv inputs, 26.8 s for Fab inputs and 6.4 s for VHH inputs. These measurements include fresh structure preparation, surface-feature extraction and prediction of residue probabilities. Fab inputs took longer because surface features are computed for the entire supplied antibody fragment, including its constant domains. Initial model loading, structure generation and visualization are excluded; response times in a web interface also depend on deployment hardware and queueing. Input sizes, component timings and measurement conditions are provided in Supplementary Section S7.

## 3. Discussion

AntiSite supports sequence-only and structure-aware paratope prediction through a shared antigen-agnostic predictor. Its modest absolute PR-AUC gains over standalone ParaSurf are accompanied by the practical benefit of supporting both input settings with shared downstream weights. Structure-aware inference additionally requires a separately paratope-trained ParaSurf encoder matched to the AntiSite checkpoint. AntiSite learns to integrate the encoder’s internal surface features before assigning probabilities. The modality-dropout ablation showed that exposure to missing structural features during training improved sequence-only PR-AUC. This flexibility allows an antibody to be analyzed first from its sequence and revisited when a structure becomes available, without retraining or switching to a different downstream predictor. The released model also demonstrated zero-shot generalization to additional antibodies and VHHs without retraining or fine-tuning.

The results support different roles for the two input settings. Sequence-only AntiSite provides a rapid first estimate without structure generation, while reliable structural input can refine the predicted binding site. Sequence-only PR-AUC was lower than Paraplume on four of five antibody evaluations, with the largest difference on AACDB. The largest gains came from combining sequence representations with molecular-surface features, which preferentially increased probabilities at contact residues. Structure-aware AntiSite also exceeded the score ensemble of sequence-only AntiSite and ParaSurf in PR-AUC across all five benchmarks. The equal-weight ensemble itself had lower reported PR-AUC than standalone ParaSurf on four of the five benchmarks. Simply averaging the final predictions therefore did not reproduce the gains obtained by learned feature integration in these comparisons. The surface-only control supports the contribution of the sequence branch beyond a newly trained head on frozen surface features. Concatenation recovered most of this improvement; attention added a modest structure-aware refinement, with no consistent sequence-only benefit (Supplementary Table S5). These results support learned feature-level fusion as the principal architectural contribution.

The matched framework comparison extends the benefit beyond the CDRs: AntiSite achieved higher PR-AUC and ROC-AUC than ParaSurf. At threshold 0.5, AntiSite selected fewer residues with higher precision and lower recall. These results distinguish improved contact ranking from the precision–recall trade-off at a particular decision threshold.

Within the tested architecture and training configuration, the PLM comparisons support using ProtT5 and ESM-2 as AntiSite’s sequence encoders. The ranking may depend on optimization and model capacity, and encoder-specific hyperparameter tuning could yield different results. The gate also learned a regional preference, assigning higher weights to ProtT5 in CDRs and to ESM-2 in frameworks without CDR/framework annotations or predefined regional rules. This shows that paratope-prediction training can produce region-dependent mixing of general protein representations. The ablations assess predictive performance under the tested settings, while the gate profiles describe the learned allocation.

Predicted antibody structures can improve paratope prediction over sequence alone, but their availability does not guarantee a performance gain. AntiSite tended to improve more when the predicted binding loops were more accurate. This pattern also held when comparing different structures of the same antibody. The most accurate predicted structures gave performance close to that obtained with experimental structures. The differences observed for ESMFold2 also show why backbone agreement alone is an incomplete description of the surface features provided to a model. Predictor confidence offers useful context when the reference structure is unknown, although the tested confidence-based switch produced only small average gains. For routine use, structures from antibody-only predictors with demonstrated benefit are reasonable optional inputs; the tested pLDDT-70 cutoff does not support a universal rule for discarding structural information.

The folding and co-folding comparisons are a further contribution of this study, providing practical evidence for choosing structural inputs for paratope prediction. In the tested cohort, providing the antigen during structure generation gave no consistent additional paratope benefit over the best antibody-only input. Direct interfaces were less accurate on average, but the highly confident subset narrowed that difference, and some model-specific subsets performed better than AntiSite with Ibex holo inputs. Co-folding therefore remains useful when an antigen epitope and a binding pose are required. AntiSite provides a paratope estimate with fewer required inputs.

Zero-shot transfer to VHHs extends this capability to antibodies lacking a light chain. This complements approaches such as AbNatiV2 that explicitly model both conventional antibodies and nanobodies[24], while testing transfer from paired-antibody training alone. The observed gains suggest that the learned contact cues are useful across these formats, although broader VHH-family coverage and VL-only single domains remain to be evaluated.

Several evaluation boundaries remain. Experimental antibody inputs retain the antigen-bound conformation and do not represent the full diversity of unbound antibodies. Similarity filtering reduces overlap with known antibody training references, but cannot guarantee independence from all upstream encoder data or antigen disjointness for antigen-aware methods. Baseline training conditions and prospective coverage also differ; these are documented in Supplementary Sections S1, S2 and S4. Related studies of affinity prediction, VH–VL pairing and VHH non-specific binding likewise show that dataset construction influences apparent generalization[20,22,23,32]. The conclusions therefore concern the evaluated cohorts and input conditions.

Paratope prediction complements antigen-side epitope prediction, exemplified by Epi4Ab[33], and affinity modeling, such as AlphaBind[34]. The use of AntiSite predictions as docking constraints or to prioritize residues for mutational analysis remains to be evaluated. Because the model is antigen-agnostic, identical antibody inputs cannot distinguish alternative antigen-dependent contact patterns. Moreover, a structural contact score does not establish energetic importance, binding affinity or the effect of a mutation; those outcomes were not evaluated here. Within this computational scope, AntiSite provides immunologists with an open tool for locating candidate antibody and nanobody binding sites, including when antigen information or experimental structures are unavailable. Its trained checkpoints enable sequence-based annotation and refinement with reliable structural input, while the folding comparisons provide evidence for choosing that input. Together, these resources support binding-interface analysis and hypothesis generation.

## 4. Materials and methods

### 4.1. Antibody inputs and reference paratopes

All benchmark evaluations followed the same antibody–antigen contact definition: an antibody residue was labeled as part of the paratope if any of its non-hydrogen atoms lay within 4.5 Å of a non-hydrogen antigen atom. Inputs comprised observed antibody variable-region residues, with the heavy chain followed by the light chain; VHH inputs contained only the heavy chain. For each antibody residue, we matched its sequence features, structural features and contact label so that all three referred to the same position. AntiSite’s structural features were computed from antibody coordinates alone. Supplementary Section S1 provides further input-processing details.

### 4.2. Benchmarks, partitions and independent evaluations

PECAN, Paragraph-expanded and MIPE retained their published partitions[8,10,11]. Processing failures were excluded without moving examples between partitions. After preprocessing, training/validation/test sizes were 195/101/152 complexes for PECAN and 624/214/216 for Paragraph-expanded. MIPE used a pool of 528 complexes split into five training/validation folds (422/106 or 423/105 complexes per fold), with a fixed test set of 63 complexes; we averaged the five models’ metrics rather than ensembling predictions. For AACDB[27], concatenated heavy/light variable-region sequences were clustered at 70% identity with CD-HIT[35], and whole clusters were assigned to training, validation or test using seed 42. AntiSite used 3,994 training and 674 validation complexes, with valid test AUC metrics for 544 complexes. Separate AntiSite checkpoints were trained on these benchmark-specific partitions.

The primary SAbDab2, VHH and predicted-structure evaluations used the released Paragraph-expanded checkpoint without retraining. Its architecture, hyperparameters and checkpoint were fixed before construction of our SAbDab2 cohort; SAbDab2 results did not inform model development or selection. SAbDab2 paired-antibody complexes came from its antigen-agnostic antibody split, version 0.2.0[28]. We retained one instance per PDB entry with a paired variable region and one contacting protein or peptide antigen. Antibodies were filtered against Paragraph-expanded and Paraplume reference sequences using an 85% identity threshold on IMGT CDR-H3 or the concatenated heavy- or light-chain CDRs. ANARCI[36] provided CDR assignments for similarity filtering, following SAbDab2’s convention; ANARCII[37] provided IMGT numbering for regional evaluation using ParaSurf’s expanded CDR boundaries. Additional baseline compatibility and training-overlap checks defined the shared test set of 178 complexes; no SAbDab2 examples were used for training or validation. The predicted-structure collection used recent complexes with the same antibody-reference filtering. The VHH evaluation retained genuine single-heavy-domain complexes, checked for overlap with AntiSite/ParaSurf training references; no VHH examples were used for training or selection. Supplementary Section S1 provides memberships, filtering stages and baseline-specific checks.

### 4.3. Protein language models and the sequence encoder

Frozen ProtT5 and ESM-2 encoders produce 1,024- and 1,280-dimensional contextual residue vectors, respectively[25,26]. Both learn sequence dependencies from general protein collections. Heavy and light sequences are concatenated for encoding; special tokens and padding are excluded from residue predictions. Embeddings are cached, and PLM parameters remain frozen throughout training.

Learned linear projections map each vector *e_i_*^(*k*)^ to a 256-dimensional representation *h_i_*^(*k*)^. A gate assigns softmax weights to the two projected vectors at each residue:

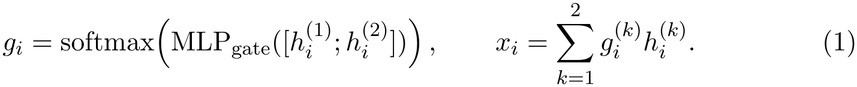

Here, *k* = 1 and *k* = 2 denote ProtT5 and ESM-2, respectively; the two weights are non-negative and sum to one at each residue. The gate is trained through the paratope-prediction losses, using only the projected PLM vectors as inputs; no CDR/framework annotations or predefined regional weights are used. A learned heavy/light chain embedding and sinusoidal position encoding are added to *x_i_*, allowing the model to distinguish chain identity and residue order. A four-layer Transformer[38] contextualizes residues across both chains, followed by layer normalization to give the *L ×* 256 representation *Z*_seq_. Padding is masked from attention. Table 4 specifies the architecture and training settings; encoder specifications and PLM comparisons are in Supplementary Sections S2–S3.

**Table 4.** AntiSite architecture and training settings. ProtT5, ESM-2 and ParaSurf are frozen; downstream layers are trained. Further implementation details are given in Supplementary Section S2.

| Component | Configuration |
|---|---|
| *Architecture* |  |
| Sequence inputs | ProtT5 (1,024 dimensions); ESM-2 650M (1,280 dimensions) |
| PLM fusion | 256-dimensional projections; learned two-way gate |
| Sequence encoder | Four chain-aware Transformer layers; eight heads; width 256 |
| Surface input | ParaSurf features; 256 dimensions per residue |
| Cross-modal attention | One block; four heads; width 256 |
| Prediction heads | Shared: 512 → 256 → 1; auxiliary: 256 → 1 (training only) |
| *Training* |  |
| Loss | Sum of masked auxiliary and shared-head BCE losses |
| Optimizer | AdamW; learning rate 10<sup>−4</sup>; weight decay 10<sup>−5</sup> |
| Batch and clipping | Eight complexes; maximum gradient norm 1.0 |
| Dropout | Network: 0.1; whole-batch structural: 0.5 |
| Model selection | Best validation structure-aware PR-AUC; patience seven; at most 100 epochs |

### 4.4. Molecular-surface features from ParaSurf

The optional structural input comprises features extracted from our previously developed ParaSurf model[9], a surface-based deep learning framework for antibody paratope prediction. DMS generates the antibody molecular surface[39]. OpenBabel provides chemical descriptors[40], and PDB2PQR assigns atomic charges and radii[41]. Surface-normal-oriented neighborhoods are voxelized on a 41 *×* 41 *×* 41 grid at 1 Å resolution. The 22 channels describe atom classes, chemical properties, charges and radii. ParaSurf’s residual CNN, Transformer and global average pooling produce a 256-dimensional vector before classification, describing the local spatial and physicochemical environment.

For residue *i*, AntiSite receives the mean of these vectors over its heavy-atom-associated surface points *A_i_*:

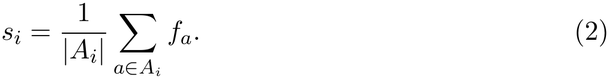

This provides structural representations rather than ParaSurf predictions; standalone ParaSurf instead uses the maximum atom-level probability for residue scoring. Missing features are zero-filled and recorded in a coverage mask. Only points needed for residue aggregation are processed.

ParaSurf is pretrained with paratope labels and remains frozen during AntiSite training; its classifier is not used inside AntiSite. Structure-aware inference therefore runs two task-trained networks, the ParaSurf encoder and the shared AntiSite predictor, while sequence-only inference runs the latter without ParaSurf. Each AntiSite checkpoint is paired with the ParaSurf extractor used to generate its training features. PECAN and Paragraph-expanded use their corresponding extractors, AACDB uses the extractor trained on the AACDB training partition, and MIPE uses a fixed fold extractor across downstream folds. The Paragraph-expanded extractor is used for all additional evaluations. Exact feature channels and checkpoint provenance are given in Supplementary Section S2.

### 4.5. Cross-modal attention and the shared prediction head

One cross-modal block uses sequence representations as queries and surface features *S* as keys and values. Four-head attention follows separate layer normalizations, with padded keys masked:

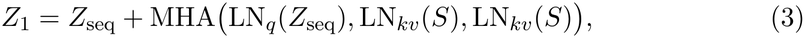

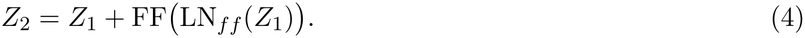

The feed-forward block uses GELU and dropout. The shared prediction head receives [*Z*_2*,i*_; *s_i_*], preserving direct access to the original surface features, and outputs a sigmoid paratope probability for every selected residue. Sequence-only inference replaces *S* with zeros while retaining the same forward pass and head. The attention block remains active, with no antibody-specific structural signal. Supplementary Figure S1 shows the implemented residual connections.

### 4.6. Training, modality dropout and ablations

During training, the complete surface tensor is zeroed for a minibatch with probability 0.5. This modality dropout[42] trains the shared parameters with and without structural input. Training minimizes the sum of two masked binary cross-entropies:

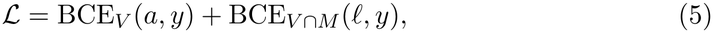

where *a* is the auxiliary-head logit computed from *Z*_seq_, *ℓ* the shared-head logit, *V* the valid-residue mask and *M* the surface-coverage mask. Each loss is averaged over eligible residues. Coverage is retained when structural features are zeroed. The auxiliary head supervises the sequence encoder during training only.

The trainable layers were optimized with AdamW[43]. Checkpoints maximized validation structure-aware PR-AUC with early stopping (Table 4). Ablations varied attention depth, head count, modality dropout and PLM composition. A surface-only head was trained on frozen ParaSurf features without sequence input or cross-modal attention. Attention-depth, surface-only and PLM comparisons used seeds 0–4; MIPE variability reflects fold-models. We also tested ESM-C 6B (ESM Cambrian), Biohub’s general protein language model[19], in place of ESM-2 while retaining ProtT5. Supplementary Section S3 reports all comparisons.

### 4.7. Baseline predictors and predicted structural inputs

The positional baseline assigns each residue its training-set contact frequency at that sequence position, following ParaSurf[9]. Each benchmark used its training partition; SAbDab2 used Paragraph-expanded training frequencies without refitting.

The benchmark comparison combines published reference results with local evaluations. Published scores retain the model configurations and evaluation conventions reported in the cited studies. On PECAN, Paragraph-expanded and MIPE, ParaSurf was re-evaluated from its released benchmark-specific weights using the same residue-level evaluation pipeline as AntiSite[9]. For AACDB, AntiSite, ParaSurf, Paraplume and ParaEM were trained on the shared training partition and evaluated on its held-out test set. SAbDab2 and VHH comparisons used released predictors on common test complexes, scored residues and contact labels. Numerical sources, processing exclusions and cohort-specific checks are detailed in Supplementary Section S1.

The score ensemble was the equal-weight mean of the per-residue probabilities from standalone ParaSurf and sequence-only AntiSite. The latter used the shared prediction head with zeroed structural features; the ensemble required no additional training.

Predicted structural inputs were compared with fixed AntiSite weights, sequences, scored residues and contact labels. ABodyBuilder3, Ibex apo/holo and Protenix-v1 provided antibody-only predictions[12–14]; Ibex holo predicts a bound-like conformation without antigen input. Boltz-2, Chai-1, Protenix-v1 and ESMFold2 received antibody and antigen sequences for complex prediction[14,17–19]. Their antibody coordinates were extracted and cropped to the matched variable region before surface processing. Model selection used predictor confidence, not experimental contacts. For direct interfaces, the minimum predicted heavy-atom distance *d_i_* to antigen was converted to *q_i_* = [1 +exp(*d_i_ −* 4.5)]*^−^*^1^, giving a 4.5-Å contact decision at 0.5. AlphaFold 3 was not evaluated because its model parameters were not available to us. Predictor settings, cutoffs and processing exceptions are detailed in Supplementary Section S4.

### 4.8. Structural accuracy, confidence and residue analyses

Predicted and experimental structures were aligned by corresponding variable-region residues and superposed on shared Fv C*α* atoms. In that common frame, we measured CDR-H3 and all-CDR backbone RMSD and reference-paratope heavy-atom RMSD. ESMFold2 was excluded from pooled backbone-error analysis because of side-chain defects. Correlations were calculated across predictions and within antibodies across alternative inputs. Confidence analyses used fixed ipTM boundaries of 0.6/0.8 and mean CDR-H3 pLDDT boundaries of 70/90 (Supplementary Section S5).

CDR annotations were not provided as inputs to AntiSite. The framework comparison used ParaSurf predictions recomputed on the same eligible antibodies. UMAP projections[44] were fitted separately for each representation; feature displacements were measured within each space against experimental inputs. Supplementary Section S6 gives region definitions and visualization settings. Molecular views used ChimeraX[45], with 100*×* predicted probabilities stored in the PDB B-factor column for coloring and antigens displayed solely for context.

### 4.9. Performance metrics and statistical comparisons

For local evaluations, including the re-evaluated ParaSurf baseline, whole-complex PR-AUC was calculated as scikit-learn average precision for each complex, then averaged across complexes. ROC-AUC uses continuous scores; F1 and Matthews correlation coefficient use threshold 0.5 in our evaluations. These metrics are also calculated per complex and averaged, weighting complexes equally. Standard deviations describe variation across complexes unless seeds or folds are specified.

Locally documented paired comparisons use two-sided Wilcoxon signed-rank tests on per-complex PR-AUC differences. Holm adjustment[29] is applied separately to the six SAbDab2 and four VHH comparisons in Figure 3 to control the family-wise error rate within each set of comparisons. A VHH sensitivity analysis averages PR-AUC within SAbDab2 antibody identifiers and repeats the four comparisons with a separate Holm correction (Supplementary Table S3B). Paired-test p-values for the predicted-structure comparisons in Table 3 are unadjusted for multiple testing and are interpreted descriptively. The separately provided AACDB paired-test summary is documented in Supplementary Section S1. Recovered structural gain is a ratio of cohort mean differences, (*P̄* _pred_ *− P̄* _seq_)*/*(*P̄* _exp_ *− P̄* _seq_), with 95% percentile intervals from 10,000 paired bootstrap resamples of complexes. Pooled structural-error and confidence summaries include repeated predictions of the same antibodies and are descriptive; within-antibody correlations provide the complementary complex-level analysis.

### 4.10. Implementation and timing

AntiSite is implemented in PyTorch. Training and inference used an NVIDIA RTX 3090. Runtime measurements used an Intel Core i7-10700K workstation with this GPU, with models resident in memory and CUDA synchronization bracketing the timed operations. Mean structure-aware inference times, including input preparation, were 13.2 s for Fv inputs, 26.8 s for Fab inputs and 6.4 s for VHH inputs. Model loading, structure generation, visualization and web-interface overhead are excluded. Supplementary Section S7 gives input sizes, component timings and measurement conditions.

## Abbreviations

AACDB: Antigen–Antibody Complex Database
BCE: Binary cross-entropy
CDR: Complementarity-determining region
CNN: Convolutional neural network
Fab: Antigen-binding fragment
Fv: Variable fragment
GELU: Gaussian error linear unit
IMGT: International ImMunoGeneTics information system
ipTM: Interface predicted template modeling score
MCC: Matthews correlation coefficient
MLP: Multilayer perceptron
PDB: Protein Data Bank
pLDDT: Predicted local distance difference test score
PLM: Protein language model
PR-AUC: Area under the precision–recall curve
RMSD: Root-mean-square deviation
ROC-AUC: Area under the receiver operating characteristic curve
SAbDab: Structural Antibody Database
SD: Standard deviation
UMAP: Uniform manifold approximation and projection
VH: Heavy-chain variable domain
VHH: Variable domain of a heavy-chain-only antibody
VL: Light-chain variable domain

## Author contributions

Angelos-Michail Papadopoulos: Conceptualization, Methodology, Software, Validation, Formal analysis, Investigation, Data curation, Visualization, Writing – original draft. Federico Alvarez: Supervision, Writing – review and editing. Petros Daras: Supervision, Project administration, Resources, Writing – review and editing.

## Declaration of generative AI use

The manuscript was drafted by the authors. OpenAI Codex (GPT-6.1) was used to assist with language polishing, including grammar correction and improvements to clarity, readability, and flow. The authors reviewed and verified all AI-assisted edits, confirmed that the tool’s terms of use were suitable for publication, and take full responsibility for the originality, accuracy, and integrity of the manuscript.

## Data and code availability

Source code, trained checkpoints and evaluation utilities are available at 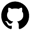 https://github.com/aggelos-michael-papadopoulos/AntiSite. Processed structures and split metadata for the five antibody benchmarks and the VHH test set are archived at 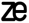 https://doi.org/10.5281/zenodo.23949886. Supplementary data provide the additional cohort memberships, per-complex comparisons and analysis summaries, with input provenance documented in Supplementary Section S1.

