## Supplementary information for "AntiSite: Unified paratope prediction for antibodies and nanobodies with or without structure"

These analyses support the benchmark comparisons and explain AntiSite's model choices. Locally calculated PR-AUC denotes average precision, except for the regional analysis in Section S6.

#### S1 Datasets and additional benchmark comparisons

The antibody evaluation covers five benchmarks. Separate models were trained for PECAN, Paragraph-expanded, MIPE and AACDB; the primary SAbDab2 and nanobody comparisons used the released Paragraph-expanded model without retraining.

##### S1.1 Training and test partitions

PECAN, Paragraph-expanded and MIPE retain their published partitions[1–3] (Table S1); MIPE's five models share one fixed test set. AACDB heavy/light sequences were clustered at 70% identity, with whole clusters assigned to training, validation or test: 3,994, 674 and 546 complexes[4].

Table S1: Published benchmark partitions and complexes retained after preprocessing.

| Benchmark | Membership | Train | Validation | Test |
|---|---|---|---|---|
| PECAN | Published | 205 | 103 | 152 |
| | Retained | 195 | 101 | 152 |
| Paragraph-expanded | Published | 651 | 217 | 218 |
| | Retained | 624 | 214 | 216 |
| MIPE | Published | 563, training/validation pool | | 63 |
| | Retained | 528, training/validation pool | | 63 |

Two cached AACDB inputs contain no contact residues, leaving 544 valid AntiSite AUC cases; ParaEM scored all 546. Restricting comparisons to 531 complexes with identical residue selections changes mean PR-AUC by at most 0.004.

##### S1.2 Independent antibody and nanobody evaluations

Similarity and compatibility filtering of the SAbDab2 antibody split[5] produced the shared 178-complex test set (Table S2). The primary ProtT5/ESM-2 model was released on 29 July 2026, before our SAbDab2 cohort was constructed on 6 October 2026. SAbDab2 results were not used to choose its architecture, hyperparameters or checkpoint.

Table S2: Filtering SAbDab2 to the shared paired-antibody test set.

| Selection step | Number retained |
|---|---|
| Original SAbDab2 antibody test structures | 3,162 |
| Paired antibodies with one contacting antigen; one complex per PDB entry | 560 |
| After excluding antibodies similar to Paragraph-expanded or Paraplume references | 185 |
| After molecular-surface preparation | 184 |
| After applying ParaDeep's chain-length limit | 181 |
| After excluding potential ParaAntiProt training overlap | 178 |

Antibodies with at least 85% identity in IMGT CDR-H3 or concatenated heavy- or light-chain CDRs were excluded against Paragraph-expanded and Paraplume references. All methods used identical antibody residues and contact labels; the ensemble equally averages sequence-only AntiSite and ParaSurf probabilities.

Table S3A provides the paired PR-AUC comparisons behind main Figure 3, with Holm adjustment separately for SAbDab2 and VHH complexes; panel B checks repeated VHH antibody identities.

Table S3: Paired PR-AUC differences: structure-aware AntiSite minus comparator. (A) Complex-level comparisons behind main Figure 3. (B) VHH sensitivity analysis after averaging within antibody identities. Differences are calculated before rounding; wins count positive differences. Two-sided Wilcoxon tests use separate Holm adjustments for the six SAbDab2, four VHH complex-level and four VHH antibody-level comparisons.

| Comparison | $n$ | Mean $\Delta$ | Wins | $p$ | $p_{\mathrm{Holm}}$ |
|---|---|---|---|---|---|
| A. Complex-level comparisons | | | | | |
| SAbDab2: AntiSite (seq) | 178 | +0.076 | 141/178 | $4.4 \times 10^{-16}$ | $1.8 \times 10^{-15}$ |
| SAbDab2: ParaSurf | 178 | +0.023 | 111/178 | $5.8 \times 10^{-4}$ | $5.8 \times 10^{-4}$ |
| SAbDab2: Paraplume | 178 | +0.068 | 127/178 | $5.5 \times 10^{-11}$ | $1.7 \times 10^{-10}$ |
| SAbDab2: ParaDeep | 178 | +0.192 | 149/178 | $6.8 \times 10^{-22}$ | $4.1 \times 10^{-21}$ |
| SAbDab2: ParaEM | 178 | +0.049 | 118/178 | $1.4 \times 10^{-6}$ | $2.8 \times 10^{-6}$ |
| SAbDab2: ParaAntiProt | 178 | +0.119 | 147/178 | $2.3 \times 10^{-20}$ | $1.1 \times 10^{-19}$ |
| VHH: ParaSurf | 140 | +0.026 | 80/140 | 0.0054 | 0.0054 |
| VHH: Paraplume | 140 | +0.122 | 116/140 | $2.9 \times 10^{-17}$ | $5.8 \times 10^{-17}$ |
| VHH: ParaDeep | 140 | +0.231 | 132/140 | $1.2 \times 10^{-22}$ | $4.7 \times 10^{-22}$ |
| VHH: AntiSite (seq) | 140 | +0.117 | 118/140 | $1.7 \times 10^{-17}$ | $5.2 \times 10^{-17}$ |
| B. VHH antibody-level sensitivity | | | | | |
| VHH: ParaSurf | 104 | +0.030 | 59/104 | 0.0117 | 0.0117 |
| VHH: Paraplume | 104 | +0.110 | 86/104 | $4.3 \times 10^{-12}$ | $8.6 \times 10^{-12}$ |
| VHH: ParaDeep | 104 | +0.238 | 98/104 | $3.3 \times 10^{-17}$ | $1.3 \times 10^{-16}$ |
| VHH: AntiSite (seq) | 104 | +0.114 | 85/104 | $1.1 \times 10^{-12}$ | $3.4 \times 10^{-12}$ |

The VHH evaluation contains 140 complexes representing 104 antibodies from SAbDab2's single-heavy-domain test split. None shared a PDB identifier or exact CDR-H3 sequence with AntiSite/ParaSurf training references. Inputs contained only the VHH; no VHH examples were used for training or model selection, and no VL-only domains were included. To account for repeated antibodies, PR-AUC was averaged within SAbDab2 antibody identifiers and the four paired tests repeated across 104 identities, with a separate Holm correction. All remained significant, including the comparison with ParaSurf ($p_{\mathrm{adj}} = 0.0117$; Table S3B).

#### S1.3 Baseline comparison scope

The benchmark comparison combines published reference results with local evaluations. Published scores retain the model configurations and evaluation conventions reported in the cited studies.

Paragraph, PECAN and Paraplume-G scores follow the experimental-structure comparison in the Paraplume study[6]. ParaEM scores on PECAN, Paragraph-expanded and MIPE are published results[7]. ParaSurf was re-evaluated on these three benchmarks from its released benchmark-specific weights using the same residue-level evaluation pipeline as AntiSite[8]. For these re-evaluations, PR-AUC is average precision calculated per complex and then averaged across complexes; F1/MCC use threshold 0.5.

SAbDab2 and VHH comparisons used released predictors on common test complexes, scored residues and contact labels. For SAbDab2, ParaEM used its released Paragraph-expanded checkpoint and ParaAntiProt used the released full-chain AntiBERTy variant trained on Parapred[7, 9]. ParaEM also receives antigen sequence; the ParaAntiProt variant differs from its published CDR-averaged PECAN evaluation.

AntiSite, ParaSurf, Paraplume and ParaEM were trained on the same AACDB partition. ParaEM's configuration was selected using validation PR-AUC alone; ParaAntiProt was not evaluated. Structure-aware AntiSite exceeded ParaEM on 433 of 544 complexes, with a mean paired PR-AUC gain of 0.085 (median 0.070; two-sided Wilcoxon signed-rank $p = 5.1 \times 10^{-52}$). The sequence-only difference was not significant (0.010, $p = 0.29$).

New positional baseline results use the same 544 AACDB and 178 SAbDab2 test cases as the corresponding comparisons, with F1/MCC threshold 0.5.

### S2 How AntiSite combines sequence and surface features

AntiSite learns to combine frozen sequence and molecular-surface features. Structure-aware inference uses the separately paratope-trained ParaSurf encoder and the shared AntiSite network. Both modes retain the same downstream weights and final prediction head.

Table S4 summarizes predictor inputs and supported inference settings.

Table S4: Antibody inputs and supported inference settings of the compared paratope predictors.

| Method | Antibody input | Sequence only | Structure aware |
|---|---|---|---|
| Sequence only | | | |
| Parapred / ParaDeep / ParaAntiProt / Paraplume | Sequence | ✓ | – |
| ParaEM[a] | Sequence | ✓ | – |
| Structure required | | | |
| Paragraph / ParaSurf | Structure | – | ✓ |
| PECAN[a] | Structure | – | ✓ |
| MIPE[a] | Sequence + structure | – | ✓ |
| Both settings | | | |
| Paraplume-G | Sequence + structure | ✓[b] | ✓ |
| AntiSite (seq-only) + ParaSurf | Sequence + structure | ✓[b] | ✓ |
| AntiSite | Sequence; optional structure | ✓ | ✓ |

[a]Antigen input is also required: sequence for ParaEM and MIPE, structure for PECAN and MIPE. [b]Sequence-only fallback uses the sequence component alone; the combined predictor requires structure.

#### S2.1 Sequence features and learned mixing

ProtT5-XL (UniRef50) and ESM-2 650M produce 1,024- and 1,280-dimensional residue features. Heavy and light sequences are concatenated, special tokens are removed, and both representations are projected to 256 dimensions. The gate learns their relative weights through paratope-prediction losses, without CDR/framework annotations.

Chain embeddings and sinusoidal positions identify chain origin and residue order. Four Transformer layers then combine information across residues and chains; the main Methods give the architectural settings.

#### S2.2 What ParaSurf contributes

ParaSurf encodes local surface geometry and chemistry[8]. DMS generates the surface; OpenBabel and PDB2PQR provide chemical descriptors, charges and radii. Surface-normal-aligned neighborhoods use a $41 \times 41 \times 41$ grid at 1 Å resolution.

The 22 channels comprise 18 chemical descriptors, two force-field charge channels and two atomic-radius channels. The bundled ParaSurf featurizer specifies AMBER and PARSE for these charge/radius pairs. Chemical descriptors cover nine atom classes, hybridization, heavy- and heteroatom degree, partial charge, hydrophobicity, aromaticity, hydrogen-bond acceptor/donor properties and ring membership. AntiSite averages ParaSurf's 256-dimensional pre-classifier features over each residue's heavy-atom-associated surface points. Missing features are zero-filled and masked.

Frozen ParaSurf extractors match the AntiSite training features: PECAN, Paragraph-expanded and AACDB use their corresponding training-partition models; MIPE uses a fixed fold-1 extractor. Additional evaluations use the Paragraph-expanded extractor. ParaSurf's classifier is unused.

#### S2.3 Combining the features in both inference modes

Sequence queries attend to surface keys and values, followed by two residual updates (Figure S1). The shared head combines the updated sequence representation with the original surface features to predict residue probabilities.

Sequence-only inference sets surface features to zero and retains the same computation. The auxiliary sequence head is used only during training; the shared-head loss retains its coverage mask when structure is dropped.

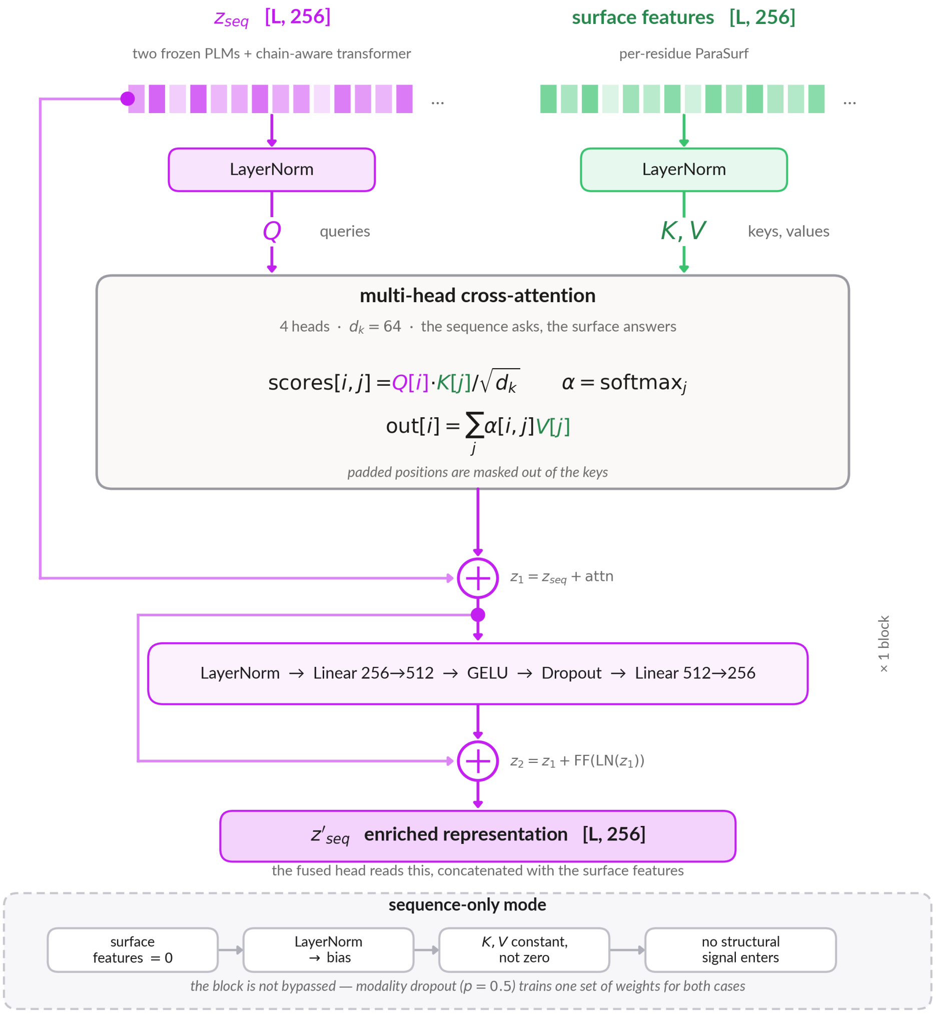


Figure S1: **How the cross-modal block combines sequence and surface features.** Sequence queries attend to ParaSurf keys and values after separate normalization. Four heads have 64 dimensions each. The first residual gives $Z_1 = Z_{\text{seq}} + \text{attention}$; the feed-forward update gives $Z_2 = Z_1 + \text{FF}(\text{LN}(Z_1))$. The shared head receives $[Z_2; S]$, of width 512. Setting $S = 0$ retains the same computation without antibody-specific surface information.

### S3 How model choices affect prediction

These ablations test attention, modality dropout and PLM choice. Checkpoints were selected by validation structure-aware PR-AUC. Five-seed ablation means need not equal the single-checkpoint scores in the main benchmark table. MIPE reports five fold-models, and single-seed sweeps are labeled separately.

#### S3.1 Sequence input, cross-modal attention and modality dropout

The surface-only control tests whether a newly trained prediction head on frozen ParaSurf features can match AntiSite without its sequence branch. It gave lower PR-AUC than both sequence–surface concatenation and full AntiSite on Paragraph-expanded and PECAN (Table S5). One attention block adds a smaller improvement over concatenation; a second gives no consistent additional benefit. Without structural-modality dropout, sequence-only PR-AUC falls to 0.496 (Table S6). The released dropout rate of 0.5 was not selected from this test-score sweep.

Table S5: Test PR-AUC for the surface-only baseline and cross-modal attention depths, mean ± SD over five seeds. The surface-only model trains a prediction head on frozen ParaSurf features without sequence input; the attention-depth variants retain both PLMs.

| Configuration | Paragraph 3D | Paragraph seq | PECAN 3D | PECAN seq |
|---|---|---|---|---|
| Surface-only | $0.789 \pm 0.0019$ | – | $0.768 \pm 0.0044$ | – |
| 0 layers | $0.812 \pm 0.001$ | $0.741 \pm 0.002$ | $0.797 \pm 0.003$ | $0.719 \pm 0.005$ |
| 1 layer (released) | $0.814 \pm 0.001$ | $0.739 \pm 0.002$ | $0.801 \pm 0.002$ | $0.721 \pm 0.003$ |
| 2 layers | $0.814 \pm 0.001$ | $0.741 \pm 0.002$ | $0.803 \pm 0.002$ | $0.719 \pm 0.005$ |

Table S6: Single-seed sensitivity analyses on Paragraph-expanded. Values are held-out test average precision. The dropout rate is the probability of zeroing all surface features in a minibatch.

**A. Cross-modal attention heads**

| Heads | 3D | Sequence only |
|---|---|---|
| 1 | 0.8128 | 0.7363 |
| 2 | 0.8146 | 0.7385 |
| 4 (released) | 0.8145 | 0.7360 |
| 8 | 0.8108 | 0.7364 |

**B. Structural-modality dropout**

| $p$ | 3D | Sequence only |
|---|---|---|
| 0.0 | 0.8138 | 0.4956 |
| 0.1 | 0.8155 | 0.7352 |
| 0.3 | 0.8169 | 0.7398 |
| 0.5 (released) | 0.8145 | 0.7360 |
| 0.7 | 0.8111 | 0.7377 |
| 0.9 | 0.8053 | 0.7445 |

#### S3.2 Choosing the protein language models

Under the common AntiSite architecture and training settings, ProtT5 plus ESM-2 gave higher PR-AUC in both modes than the antibody-specific AbLang2/IgT5 pair and the four-encoder stack on PECAN and Paragraph-expanded[10, 11]; the general pair and four-encoder stack performed similarly on MIPE (Table S7). Table S8 tests removing each encoder from the four-model stack, with separate training for each configuration.

Table S7: Test PR-AUC by PLM combination, mean ± SD over five seeds. MIPE values are means across five fold-models.

| Stack | Paragraph 3D | Paragraph seq | PECAN 3D | PECAN seq |
|---|---|---|---|---|
| Four PLMs | $0.810 \pm 0.001$ | $0.730 \pm 0.002$ | $0.794 \pm 0.002$ | $0.715 \pm 0.003$ |
| ProtT5 + ESM-2 | $0.814 \pm 0.001$ | $0.739 \pm 0.002$ | $0.801 \pm 0.002$ | $0.721 \pm 0.003$ |
| AbLang2 + IgT5 | $0.807 \pm 0.001$ | $0.715 \pm 0.006$ | $0.790 \pm 0.001$ | $0.685 \pm 0.002$ |

MIPE fold-model averages: four encoders, 3D 0.790 / sequence only 0.716; general pair, 3D 0.789 / sequence only 0.718. Fold-models share the 63-complex test set.

Table S8: Five-seed test PR-AUC (mean ± SD) after removing each encoder from the AbLang2/ProtT5/ESM-2/IgT5 stack.

| Configuration | Paragraph 3D | Paragraph seq | PECAN 3D | PECAN seq |
|---|---|---|---|---|
| Full four-model stack | $0.810 \pm 0.001$ | $0.730 \pm 0.002$ | $0.794 \pm 0.002$ | $0.715 \pm 0.003$ |
| Drop AbLang2 | $0.811 \pm 0.002$ | $0.733 \pm 0.002$ | $0.797 \pm 0.002$ | $0.716 \pm 0.003$ |
| Drop IgT5 | $0.812 \pm 0.000$ | $0.735 \pm 0.004$ | $0.798 \pm 0.003$ | $0.718 \pm 0.004$ |
| Drop ProtT5 | $0.810 \pm 0.001$ | $0.730 \pm 0.004$ | $0.792 \pm 0.003$ | $0.707 \pm 0.005$ |
| Drop ESM-2 | $0.811 \pm 0.002$ | $0.726 \pm 0.001$ | $0.794 \pm 0.002$ | $0.710 \pm 0.003$ |

In a post hoc comparison, we tested ESM-C 6B (ESM Cambrian), a general protein language model from Biohub with approximately six billion parameters[12], as a replacement for ESM-2 while retaining ProtT5. With five matched seeds, this replacement improved sequence-only PR-AUC but gave no significant structure-aware gain (Table S9). The released model retains ProtT5 and ESM-2.

Table S9: Post hoc five-seed comparison of ProtT5 paired with ESM-2 or ESM-C 6B: test PR-AUC (mean ± SD). Paired p-values use per-complex PR-AUC averaged across seeds.

| Evaluation | Mode | $n$ | ProtT5 + ESM-2 | ProtT5 + ESM-C | $p$ |
|---|---|---|---|---|---|
| Paragraph test | 3D | 216 | $0.8137 \pm 0.0026$ | $0.8161 \pm 0.0010$ | 0.2595 |
| Paragraph test | seq | 216 | $0.7369 \pm 0.0049$ | $0.7467 \pm 0.0044$ | 0.0010 |
| SAbDab2 (earlier) | 3D | 184 | $0.7486 \pm 0.0010$ | $0.7444 \pm 0.0027$ | 0.3667 |
| SAbDab2 (earlier) | seq | 184 | $0.6724 \pm 0.0042$ | $0.6806 \pm 0.0060$ | 0.0197 |

Both variants were trained on Paragraph-expanded training data and selected by its validation PR-AUC. This comparison followed the release of the primary ProtT5/ESM-2 model and did not guide its selection. The 184 SAbDab2 cases precede ParaDeep's chain-length and ParaAntiProt's training-overlap exclusions (Table S2); the main comparison uses the resulting 178-case subset.

### S4 Predicted antibody structures and evaluation scope

These comparisons test whether predicted structures help AntiSite. The released weights remain fixed, and AntiSite receives only antibody sequence and coordinates, including when the coordinates come from antigen-aware co-folding. Paired-test p-values in these analyses are unadjusted for multiple testing and reported descriptively.

The 87 complexes were released on or after 1 February 2024 and filtered against antibody training references. Table S10A lists predictor cutoffs; panel B gives the paired two-sided Wilcoxon signed-rank tests against sequence-only predictions for each structural input. When multiple predictions were available, selection used predictor confidence. Chai-1 provided 86 structures and ESMFold2 80, with comparisons restricted to matched available cases.

Table S10: Predictor inputs, documented structural-training cutoffs and paired tests of predicted structural inputs. Cutoffs do not establish independence from all upstream pretraining data.

**A. Predictor inputs and training cutoffs**

| Predictor | Input setting | Structural-training cutoff |
|---|---|---|
| ABodyBuilder3 | Antibody only | January 2024 |
| Ibex apo / holo | Antibody only | Not published |
| Protenix-v1 | Antibody only and co-folding | 30 September 2021 |
| Boltz-2 | Antibody–antigen co-folding | 1 June 2023 |
| Chai-1 | Antibody–antigen co-folding | 12 January 2021 |
| ESMFold2 | Antibody–antigen co-folding | September 2021 |

Co-folding uses the antigen during structure generation; AntiSite's surface features use only the predicted antibody coordinates.

**B. Paired tests against sequence-only predictions**

| Structural input | Folding setting | $n$ | Unadjusted $p$ |
|---|---|---|---|
| Protenix-v1 | Antibody only | 87 | 0.00209 |
| Ibex (apo) | Antibody only | 87 | $6.23 \times 10^{-5}$ |
| Ibex (holo) | Antibody only | 87 | $3.24 \times 10^{-5}$ |
| ABodyBuilder3 | Antibody only | 87 | 0.528 |
| Boltz-2 | Co-folding | 86 | $1.38 \times 10^{-5}$ |
| Chai-1 | Co-folding | 86 | 0.0354 |
| Protenix-v1 | Co-folding | 86 | $1.61 \times 10^{-5}$ |
| ESMFold2 | Co-folding | 80 | 0.691 |

Two-sided paired Wilcoxon signed-rank tests compare per-complex PR-AUC with sequence-only predictions on the same cohort. P-values are unadjusted for multiple testing and reported descriptively.

Because Ibex has no published training cutoff, Table S11 also evaluates 36 complexes released after its 11 July 2025 weight release. Predicted-input gain intervals include zero in this subset.

Table S11: Matched post-Ibex-release subset ($n = 36$).

| Input | PR-AUC | Gain recovered | 95% interval (%) | $p$ vs seq |
|---|---|---|---|---|
| Sequence only | 0.734 | – | – | – |
| Protenix-v1 | 0.741 | 10% | −40 to 42 | 0.49 |
| Ibex (apo) | 0.753 | 27% | −5 to 57 | 0.16 |
| Ibex (holo) | 0.748 | 20% | −14 to 53 | 0.23 |
| ABodyBuilder3 | 0.723 | -17% | −74 to 22 | 0.46 |
| Experimental structure | 0.803 | 100% | – | $2.1 \times 10^{-6}$ |

Unrefined IgFold predictions lacked complete side chains and could not be processed by ParaSurf[13]. The evaluated ESMFold2 outputs contained side-chain clashes that compressed AntiSite probabilities. They were retained as measured, without experimental side-chain replacement, but excluded from pooled backbone-error analysis because backbone RMSD does not capture these defects.

### S5 Structural accuracy and predictor confidence

We examined whether structural accuracy explains when predicted structures help, and whether predictor confidence offers guidance without an experimental reference.

#### S5.1 Measuring structural error

Lower RMSD indicates closer agreement with the experimental structure. Structures were superposed on shared Fv C$\alpha$ atoms, without realigning individual loops. Backbone errors use matched N/C$\alpha$/C/O atoms; paratope errors use matched heavy atoms in contact residues (Table S12).

Table S12: Median structural errors (Å) after shared Fv superposition, and mean AntiSite PR-AUC gain over sequence-only inference. Each row uses the predictor's available cases.

| Input | $n$ | Fv C$\alpha$ | H3 backbone | All CDRs | Contact atoms | $\Delta$ PR |
|---|---|---|---|---|---|---|
| Protenix-v1 (Ab only) | 87 | 0.93 | 1.83 | 1.51 | 2.50 | +0.031 |
| Ibex (apo) | 87 | 1.00 | 2.15 | 1.50 | 2.63 | +0.035 |
| Ibex (holo) | 87 | 0.87 | 1.78 | 1.29 | 2.22 | +0.038 |
| ABodyBuilder3 | 87 | 1.27 | 3.15 | 2.01 | 3.43 | -0.008 |
| Boltz-2 | 87 | 1.00 | 2.42 | 1.57 | 3.02 | +0.033 |
| Chai-1 | 86 | 1.14 | 2.58 | 1.68 | 2.79 | +0.019 |
| Protenix-v1 (co-fold) | 87 | 0.91 | 1.95 | 1.39 | 2.47 | +0.039 |
| ESMFold2† | 80 | 1.15 | 2.17 | 1.83 | 2.86 | -0.001 |

†ESMFold2 is excluded from pooled backbone analysis because of its side-chain defects.

Comparing alternative structures of the same antibody also linked higher error to lower PR-AUC gain across all four error measures (Table S13).

Table S13: Structural error versus PR-AUC gain across alternative predictions for each of 87 antibodies. Signed-rank tests compare these antibody-level correlations with zero.

| Error measure | Mean $\rho$ | Negative correlations | $p$ |
|---|---|---|---|
| Fv C$\alpha$ RMSD | -0.375 | 83% | $5.6 \times 10^{-10}$ |
| CDR-H3 backbone RMSD | -0.383 | 78% | $1.9 \times 10^{-9}$ |
| All-CDR backbone RMSD | -0.397 | 77% | $9.8 \times 10^{-10}$ |
| Paratope all-atom RMSD | -0.369 | 78% | $1.2 \times 10^{-8}$ |

#### S5.2 Using predictor confidence

Higher interface confidence (ipTM) narrows the difference between direct co-folded interfaces and AntiSite with antibody-only Ibex holo input; some high-confidence subsets favor direct interfaces (Table S14).

Table S14: Mean PR-AUC for direct interfaces versus AntiSite with antibody-only Ibex holo input. Panel A pools Boltz-2, Chai-1 and Protenix-v1, including repeated antibodies; tests are descriptive. Panel B uses predictor-specific ipTM $\geq 0.8$ subsets, including ESMFold2.

**A. Pooled confidence bins**

| ipTM bin | Predictions | Direct interface | AntiSite + Ibex holo | $p$ |
|---|---|---|---|---|
| $< 0.6$ | 63 | 0.613 | 0.711 | $1.1 \times 10^{-6}$ |
| 0.6–0.8 | 147 | 0.677 | 0.738 | $3.7 \times 10^{-5}$ |
| $\geq 0.8$ | 50 | 0.818 | 0.826 | 0.87 |

**B. Model-specific high-confidence subsets**

| Predictor | $n$ | Direct interface | AntiSite + Ibex holo | $p$ |
|---|---|---|---|---|
| Boltz-2 | 23 | 0.747 | 0.837 | 0.0075 |
| Chai-1 | 9 | 0.887 | 0.859 | 0.43 |
| Protenix-v1 | 18 | 0.874 | 0.795 | 0.027 |
| ESMFold2 | 10 | 0.913 | 0.779 | 0.0098 |

Higher CDR-H3 confidence (pLDDT) is associated with larger structural-input gains, but switching to sequence-only inference below 70 changes mean performance little (Table S15). These thresholds are exploratory and are not validated selection rules.

Table S15: CDR-H3 confidence and structural-input gain. Panel A pools 347 predictions from Protenix-v1 (antibody-only/co-folding), Boltz-2 and Chai-1; intervals resample predictions, including repeated antibodies. Panel B adds ESMFold2 and uses sequence-only inference below pLDDT 70. Changes are calculated before rounding.

**A. Confidence bins**

| CDR-H3 pLDDT | $n$ | Mean gain | 95% interval | Median RMSD (Å) |
|---|---|---|---|---|
| $< 70$ | 42 | −0.008 | −0.031 to +0.015 | 4.76 |
| 70–90 | 218 | +0.025 | +0.014 to +0.036 | 2.43 |
| $\geq 90$ | 87 | +0.062 | +0.047 to +0.077 | 1.03 |

**B. Fixed confidence rule**

| Predictor | $n$ | Always structure | Confidence rule | Change |
|---|---|---|---|---|
| Protenix-v1 (Ab only) | 87 | 0.741 | 0.741 | +0.000 |
| Boltz-2 | 87 | 0.743 | 0.739 | −0.004 |
| Chai-1 | 86 | 0.728 | 0.737 | +0.009 |
| Protenix-v1 (co-fold) | 87 | 0.749 | 0.748 | −0.001 |
| ESMFold2 | 80 | 0.706 | 0.712 | +0.007 |

### S6 Antibody regions and residue-level plots

Regional comparisons distinguish performance on binding loops and framework residues; the residue-level plots help interpret the learned features.

#### S6.1 CDR and framework predictions

Structure-aware AntiSite exceeds recomputed ParaSurf on both framework ranking metrics and improves over sequence-only AntiSite in all six CDRs and the framework (Table S16).

The regional analysis follows ParaSurf's expanded IMGT definition[8]: CDR1 25–40, CDR2 54–67 and CDR3 103–119. Regional PR-AUC uses trapezoidal integration, so it differs from whole-complex average precision. Published ParaSurf CDR values remain included for context.

Table S16: Regional performance on Paragraph-expanded, using expanded IMGT boundaries and trapezoidal PR-AUC. Only antibodies with both contacts and non-contacts contribute. Dagger rows are published ParaSurf CDR scores; framework ParaSurf was recomputed on AntiSite's cohort. F1/MCC use threshold 0.5.

| Method | PR-AUC | ROC-AUC | F1 | MCC |
|---|---|---|---|---|
| L-CDR1 ($n = 189$) | | | | |
| ParaSurf† | 0.885 | 0.965 | 0.692 | 0.669 |
| Paraplume | 0.747 | 0.879 | 0.583 | 0.515 |
| AntiSite (sequence only) | 0.840 | 0.933 | 0.683 | 0.631 |
| AntiSite (3D) | 0.884 | 0.955 | 0.716 | 0.674 |
| L-CDR2 ($n = 109$) | | | | |
| ParaSurf† | 0.772 | 0.932 | 0.535 | 0.509 |
| Paraplume | 0.796 | 0.853 | 0.537 | 0.450 |
| AntiSite (sequence only) | 0.809 | 0.903 | 0.393 | 0.365 |
| AntiSite (3D) | 0.863 | 0.930 | 0.535 | 0.489 |
| L-CDR3 ($n = 189$) | | | | |
| ParaSurf† | 0.910 | 0.989 | 0.617 | 0.619 |
| Paraplume | 0.778 | 0.871 | 0.693 | 0.611 |
| AntiSite (sequence only) | 0.862 | 0.956 | 0.731 | 0.689 |
| AntiSite (3D) | 0.921 | 0.973 | 0.783 | 0.750 |
| H-CDR1 ($n = 191$) | | | | |
| ParaSurf† | 0.868 | 0.964 | 0.681 | 0.652 |
| Paraplume | 0.874 | 0.952 | 0.739 | 0.690 |
| AntiSite (sequence only) | 0.837 | 0.930 | 0.584 | 0.533 |
| AntiSite (3D) | 0.875 | 0.951 | 0.711 | 0.670 |
| H-CDR2 ($n = 197$) | | | | |
| ParaSurf† | 0.896 | 0.963 | 0.748 | 0.710 |
| Paraplume | 0.793 | 0.861 | 0.709 | 0.552 |
| AntiSite (sequence only) | 0.853 | 0.907 | 0.715 | 0.589 |
| AntiSite (3D) | 0.893 | 0.943 | 0.772 | 0.666 |
| H-CDR3 ($n = 211$) | | | | |
| ParaSurf† | 0.895 | 0.959 | 0.759 | 0.709 |
| Paraplume | 0.625 | 0.731 | 0.567 | 0.393 |
| AntiSite (sequence only) | 0.824 | 0.938 | 0.735 | 0.657 |
| AntiSite (3D) | 0.902 | 0.965 | 0.810 | 0.757 |
| Framework ($n = 143$) | | | | |
| ParaSurf (recomputed) | 0.546 | 0.968 | 0.402 | 0.414 |
| Paraplume | 0.340 | 0.844 | 0.171 | 0.178 |
| AntiSite (sequence only) | 0.507 | 0.965 | 0.148 | 0.156 |
| AntiSite (3D) | 0.616 | 0.974 | 0.227 | 0.234 |

At threshold 0.5, AntiSite makes fewer framework predictions with higher precision and lower false-positive rate, while ParaSurf recovers more contacts and retains higher F1/MCC (Table S17). Framework contacts are rare (about 1.2%), which should be considered when interpreting regional PR-AUC.

Table S17: Pooled framework predictions at threshold 0.5; FPR denotes false-positive rate. These complement the per-antibody averages in Table S16.

| Model | Recall | FPR | Precision | Predicted positives | Contacts recovered |
|---|---|---|---|---|---|
| ParaSurf | 0.475 | 0.0099 | 0.358 | 508 | 182 |
| AntiSite (3D) | 0.227 | 0.0009 | 0.744 | 117 | 87 |

#### S6.2 Reading the residue-level plots

Main Figure 2 uses experimental antibody coordinates and the released paired-antibody-trained checkpoint. Each format includes two high-performing examples and one near its cohort median; these illustrations were not randomly sampled. Probabilities multiplied by 100 were written to the PDB B-factor column for ChimeraX coloring. Antigens retain their experimental positions for context only, and views are independently scaled.

Main Figures 4 and 8 align residues by IMGT segment and resample to median segment lengths. Figure 4 contact curves are shown only where at least five antibodies contribute a contact residue. Figure 8 heavy- and light-chain heatmap rows are sorted separately by mean ProtT5 weight; its gate distributions cover 15,589 framework and 4,448 CDR residues. CDR annotations are used only for analysis.

Figure 9 uses cosine UMAP with 25 neighbors, minimum distance 0.25 and seed 0[14]. Columns are fitted independently with identical settings; both colorings within a column share coordinates. CDR categories are applied after fitting and are not model inputs; chain identity and position are explicit inputs. The sequence-encoder column precedes cross-modal attention and is not the complete sequence-only inference path, which also runs attention with zero structural features. The prediction head additionally receives the original surface features. These maps describe intermediate features, not predictive accuracy.

Figure 6 samples 25 matched residues per structure, using cosine UMAP with 30 neighbors, minimum distance 0.3 and seed 0. Its distances average matched-residue differences from the experimental reference within each feature space; scales cannot be compared between spaces.

### S7 Prediction time and surface-processing cost

Table S18 summarizes prediction times on an NVIDIA RTX 3090/Intel Core i7-10700K workstation with all models loaded and CUDA-synchronized timers. Sequence-only times include PLM inference. Structure-aware times also include fresh structure preparation and surface-feature extraction. Model loading, structure generation, visualization and web-interface overhead are excluded.

Table S18: Inference time by input type. Times are means ± SD in seconds across inputs; residue ranges describe the supplied structures.

| Input or mode | Fv ($n = 6$) | Fab ($n = 9$) | VHH ($n = 3$) |
|---|---|---|---|
| Observed residues | 224–237 | 421–439 | 116–129 |
| Sequence-only prediction | $0.088 \pm 0.004$ | $0.103 \pm 0.001$ | $0.072 \pm 0.007$ |
| Structure-aware prediction | $13.2 \pm 1.0$ | $26.8 \pm 2.2$ | $6.4 \pm 0.3$ |

Fab inputs take longer because ParaSurf processes constant domains as well as variable domains. Most additional computation comes from voxelization and the surface CNN. Per-input measurements and component timings are provided in the supporting data.
